# Electrophysiological Mechanisms of Psychedelic Drugs: A Systematic Review

**DOI:** 10.1101/2025.07.05.663289

**Authors:** Javier Hidalgo Jiménez, Kristjan Kaup, Jaan Aru

## Abstract

Serotonergic psychedelics are known for their profound effects on consciousness and are gaining renewed interest as potential psychiatric treatments. These advances underscore the need to clarify the mechanisms of action of these compounds. This systematic review compiles and critically evaluates 23 *in vitro* and 26 *in vivo* electrophysiological studies on psychedelic compounds, with an emphasis on layer 5 pyramidal neurons in the prefrontal cortex, where 5-HT_2A_ receptors are densely expressed. Our findings reveal that psychedelics exert complex, heterogeneous effects on neuronal excitability, synaptic transmission, and local oscillations. These results challenge the simplified view that psychedelics uniformly increase cortical excitability. Instead, they modulate both excitatory and inhibitory processes in a cell-type- and compartment-specific manner, with evidence for biphasic, dose-dependent, and context-sensitive responses. Activation of 5-HT_2A_ receptors leads to intricate calcium signaling, downregulating excitatory currents and firing rates in many neurons, while enhancing glutamate release and activating a subset of projection fibers. Modulation of presynaptic and extrasynaptic GluN2B-containing NMDA receptors appears central to these effects, and some indirect evidence supports the involvement of intracellular 5-HT_2A_ receptors. These insights prompt a reassessment of prevailing models of psychedelic action and underscore the value of incorporating electrophysiological data into psychedelic neuropharmacology.

## 1. INTRODUCTION

Psychedelics are a family of tryptamines (e.g., psilocybin), ergolines (e.g., lysergic acid diethylamide, LSD), and phenylethylamines (e.g., 2,5-dimethoxy-4-iodoamphetamine, DOI) that profoundly alter human consciousness. Plant and fungal species containing psychedelic alkaloids grow on all five continents, and evidence of their use dates back to Neolithic cave paintings (Escohotado, 2005). Their relevance to archaic religions is certain among pre-Columbian civilizations of Central America and the Andes, and possible in the Vedic culture. They were also likely used in the Eleusinian Mysteries for nearly 2,000 years, rites attended by figures such as Plato, Aristotle, Cicero and Marcus Aurelius (Escohotado, 2005; Nichols, 2016). The effects include sensory alterations such as synesthesia and hallucinations, distortions in the perception of space and time, distortions of identity (ego dissolution or ego death), and ecstatic experiences (Nichols, 2016).

Recently, psychiatry has become interested in these drugs. Psychedelics have shown promising results in clinical trials, emerging as potential treatments for psychiatric conditions with high prevalence and morbidity, such as major depression and anxiety (Davis et al., 2021; dos Santos et al., 2018; Goodwin et al., 2022; Muttoni et al., 2019; Raison et al., 2023; Schimmers et al., 2022; von Rotz et al., 2023). Computational neuroscientists have also become interested in modeling the effects of psychedelic drugs on brain dynamics (Deco et al., 2018; Herzog et al., 2023; Irrmischer et al., 2025; Kringelbach et al., 2020; Luppi et al., 2021), in an attempt to better understand brain states.

Psychedelics are partial agonists at the serotonin 2A receptor (5-HT_2A_). 5-HT_2A_ is a G protein-coupled receptor (GPCR). Through Gɑ_q/11_-mediated signaling, it activates the release of Ca^2+^ from intracellular stores. Ca^2+^ ions activate protein kinases like protein kinase C (PKC) or the Ca^2+^/calmodulin-dependent protein kinase II (CaM-KII). Protein kinases phosphorylate ionic channels in the cell, modulating excitability. Activation of 5-HT_2A_ has been shown to increase neural excitability (Aghajanian & Marek, 1999). Additionally, psychedelics promote the release of glutamate (Wojtas et al., 2022), further contributing to neuronal excitation.

The receptor is expressed in various cell types across the brain, including both excitatory and inhibitory neurons, as well as glia. However, it is most abundant in the apical dendrites of layer 5 pyramidal (L5p) neurons (Chiu et al., 2023; Weber & Andrade, 2010). Because L5p cells occupy a central position in cortical computations and are highly interconnected with the thalamus, they are considered key to brain function and consciousness (Aru et al., 2020).

Most current approaches to understanding the action of psychedelics rely on resting-state neuroimaging, a field that has seen tremendous advancements. The sophistication of fMRI and the involvement of computational neuroscientists have revolutionized psychedelic science. However, due to the inherent limitations of these methods, it is crucial to validate and support their findings with more precise physiological techniques, such as electrophysiology. Resting-state fMRI studies, for instance, often interpret increased functional connectivity as heightened neuronal activity (Ghaw et al., 2024). Yet, several studies in humans and rodents have reported a generalized decrease in BOLD signal across the brain during psychedelic states (Carhart-Harris et al., 2012; Ghaw et al., 2024; Riga et al., 2014), despite increased between-network functional connectivity. Since BOLD signal changes can reflect alterations in both inhibitory and excitatory processes, fMRI alone cannot fully disentangle these mechanisms. Electrophysiology, which directly measures neuronal firing rates, oscillatory activity, and synaptic transmission, provides the temporal and physiological precision necessary to clarify how psychedelics modulate neural function. Given these advantages, revisiting electrophysiological research (an area that thrived before the modern "renaissance" of psychedelic science) is essential for complementing and refining insights derived from neuroimaging (Smausz et al., 2022).

Several popular reviews of psychedelic activity overemphasize the excitatory effects of psychedelics (e.g., van Elk & Yaden, 2022; Vollenweider & Preller, 2020), leaving an impression that psychedelics have a neat excitatory action in neurons and circuits. Moreover, several leading hypotheses about the mode of action of psychedelics, often based on neuroimaging, rely on their ability to increase neural activity. For instance, the Relaxed Beliefs Under Psychedelics (REBUS) hypothesis proposes that increased excitability of L5p neurons dysregulates higher-order brain functions, leading to decreased top-down control (Carhart-Harris et al., 2014; Carhart-Harris & Friston, 2019). Similarly, the Cortico-Striato-Thalamo-Cortical (CSTC) loop hypothesis, inspired by classic models of schizophrenia, posits that psychedelic hallucinations arise from heightened excitability in L5p neurons, reducing thalamic inhibition of sensory input (Vollenweider & Geyer, 2001; Vollenweider & Preller, 2020).

Despite some work on psychedelics showing increased neural excitability *in vitro* and *in vivo* (Araneda & Andrade, 1991; De Gregorio et al., 2021; Shao et al., 2025), there is considerable evidence suggesting a significantly more complex picture. For instance, LSD and DOI can reduce neural firing across both excitatory and inhibitory cells in a wide range of cortical and subcortical regions in awake animals (Brys et al., 2023). Moreover, *in vivo* studies show that DOI, which lacks inhibitory 5-HT_1A_ agonist activity, reliably inhibits L5p firing in a dose-dependent manner when applied locally (Ashby et al., 1989; Ashby & Wang, 1990; Bambico et al., 2010; Bergqvist et al., 1999; El Mansari & Blier, 1997, 2005; Rueter & Blier, 1999; Reuter et al., 2000). These often-overlooked findings underscore the complexity of psychedelic effects on neuronal activity and call for a reassessment of some of the most prevailing concepts in this field of research.

Nevertheless, accessing the literature about electrophysiological research on psychedelics remains challenging. Many results are buried within studies focused on serotonin and its receptors, rather than psychedelics specifically. Additionally, some findings appear contradictory and only make sense after carefully analyzing experimental procedures and comparing work from different authors.

Here we present a systematic review dedicated to the electrophysiological effects of psychedelic drugs. We cover both *in vitro* experiments using cortical slices and *in vivo* studies.

Since 5-HT_2A_ receptors are predominantly expressed in L5p neurons we include only *in vitro* studies that examine pyramidal neurons from layers 5 and 6, as these layers are often mixed in experimental designs. Additionally, because the prefrontal cortex (PFC) is highly enriched in 5-HT_2A_ receptors and is involved in numerous cognitive functions, it has been a primary focus in psychedelic research. Therefore, we include only *in vivo* experiments that record from at least one area of the frontal cortex. Most research has been conducted with DOI, given its relative accessibility and specificity for 5-HT_2A/C_ receptors. Other psychedelic phenylethylamines included in this review are DOB, a bromine-substituted congener of DOI; and TCB-2 and 25CN-NBOH (NBOH), which exhibit high potency and selectivity for the 5-HT_2A_ receptor. We also examine LSD, which has been extensively studied in clinical psychedelic research, as well as psilocin and psilocybin, the active compounds in psychedelic mushrooms that have gained significant interest in clinical applications. Additionally, we include DMT and 5-MeO-DMT, two short-acting psychedelics, along with 4-OH-DiPT, a psilocin derivative.

The first part of the Results section will summarize *in vitro* research. We will begin by presenting the effects of psychedelics on passive membrane properties, such as resting membrane potential (RMP) and input resistance (Ri). Next, we will describe their postsynaptic effects, detailing the complex modulation of excitatory postsynaptic potentials (EPSPs) and currents (EPSCs), their underlying mechanisms, and their peculiar effects on asynchronous EPSCs. Finally, we will examine how psychedelics influence action potential spikes.

The second part of the Results section will focus on *in vivo* experiments. We will start by analyzing the effects of local microiontophoretic drug application on neuronal firing. Then, we will explore the effects of systemic drug administration in anesthetized animals, addressing both neuronal firing and oscillatory activity. Lastly, we will review *in vivo* experiments in awake animals, covering firing distribution, firing activity, and oscillatory dynamics.

Within sections, the experiments will be addressed in chronological order.

In the discussion, we propose a cohesive framework, inspired by Dendritic Integration Theory (Bachman et al., 2020), to elucidate the electrophysiological effects of psychedelics. We also introduce a unified molecular mechanism of synaptic modulation by psychedelic compounds and examine the role of 5-HT_2A_ receptor-mediated neural inhibition in mediating the effects of these substances. Finally, we situate our hypothesis within the wider context of psychedelic functional and behavioral research, while addressing the study’s limitations and outlining future directions.

## 2. METHODS

Data for this systematic review were collected following the systematic reviews and meta-analyses guidelines (Preferred Reporting Items for Systematic Reviews and Meta-Analyses [PRISMA]). The review was not registered.

### 2.1 Search

PubMed was used to identify relevant articles, with no restrictions on publication dates. Both peer-reviewed articles and preprints were included (see Tables S1 and S2 for details on non-peer-reviewed papers). The final search was performed on 05.12.2025. The following search terms were employed: : ("psychedelic*" OR "hallucinogen*" OR "LSD" OR "psilocybin" OR "psilocin" OR "ayahuasca" OR "DMT" OR "dimethyltryptamine" OR "N,N-dimethyltryptamine" OR "mescaline" OR "DOI" OR "2,5-Dimethoxy-4-iodoamphetamine" OR "DOM" OR "2,5-Dimethoxy-4-methylamphetamine" OR "DOB" OR "2,5-Dimethoxy-4-bromoamphetamine") AND ("electrophysiolog*" OR "neurophysiolog*" OR "cortical slice*" OR "brain slice*" OR "field potential*" OR "neuronal activity" OR "multi-unit recording*" OR "local field potential*" OR "spike train*" OR "neuronal oscillations" OR "patch clamp*" OR "single unit extracellular recording*" OR "microelectrode intracellular recording*") AND ("prefrontal cortex" OR "PFC" OR "medial prefrontal cortex" OR "mPFC" OR "ventromedial prefrontal cortex" OR "vmPFC" OR "dorsolateral prefrontal cortex" OR "dlPFC" OR "orbitofrontal cortex" OR "OFC" OR "pyramidal neuron" OR "pyramidal cell" OR "pyramidal neurons" OR "pyramidal cells") AND ("journal article"[Publication Type] OR "preprint"[Publication Type]) NOT review[Publication Type]. The literature search tagged both the title and the abstract. An additional search was conducted in Google Scholar to ensure comprehensive coverage of relevant literature.

### 2.2 Eligibility criteria

Studies were excluded if they met any of the following conditions:

1. They were not original research (e.g., reviews, commentaries, or meta-analyses);
2. The serotonergic agonists used do not produce in animals behavioral responses suggestive of a hallucinogenic effect in humans;
3. They did not assess electrophysiological properties of single layer 5 pyramidal neurons in vitro (such as passive membrane properties, postsynaptic currents or potentials, or action potentials);
4. They did not involve in vivo intracerebral electrophysiological recordings (e.g., single-unit activity, multi-unit activity, or local field potentials);
5. They did not examine at least one region within the frontal cortex.

### 2.3 Data extraction

After removing duplicates, all remaining articles underwent title and abstract screening. Studies that did not fulfill the eligibility criteria were excluded. Full-text assessments were conducted to determine eligibility.

For each *in vitro* included study, the following data were extracted: (1) specie; (2) recorded region; (3) specific psychedelic compound and (4) used concentration; (5) co-treatments and their concentration; effects on (6) passive membrane properties, (7) postsynaptic activity, and (8) action potential parameters; (9) other relevant findings; and (10) peer-review status.

For each *in vivo* study, the following data were extracted: (1) specie; (2) recorded region; (3) the specific psychedelic compound and (4) administered dose; (5) administration route; (6) anesthesia and co-treatments; effects on (7) neural firing (single unit or multi-unit activity) and (8) local field potentials; (9) other relevant findings; and (10) peer-review status.

Given the difficulty of including the exact number of units recorded for each result, the sample size of the experiments was not systematically extracted.

### 2.4 Identified studies

A flow diagram illustrating the different phases of the systematic review is presented in Figure 1.

**Figure 1:**
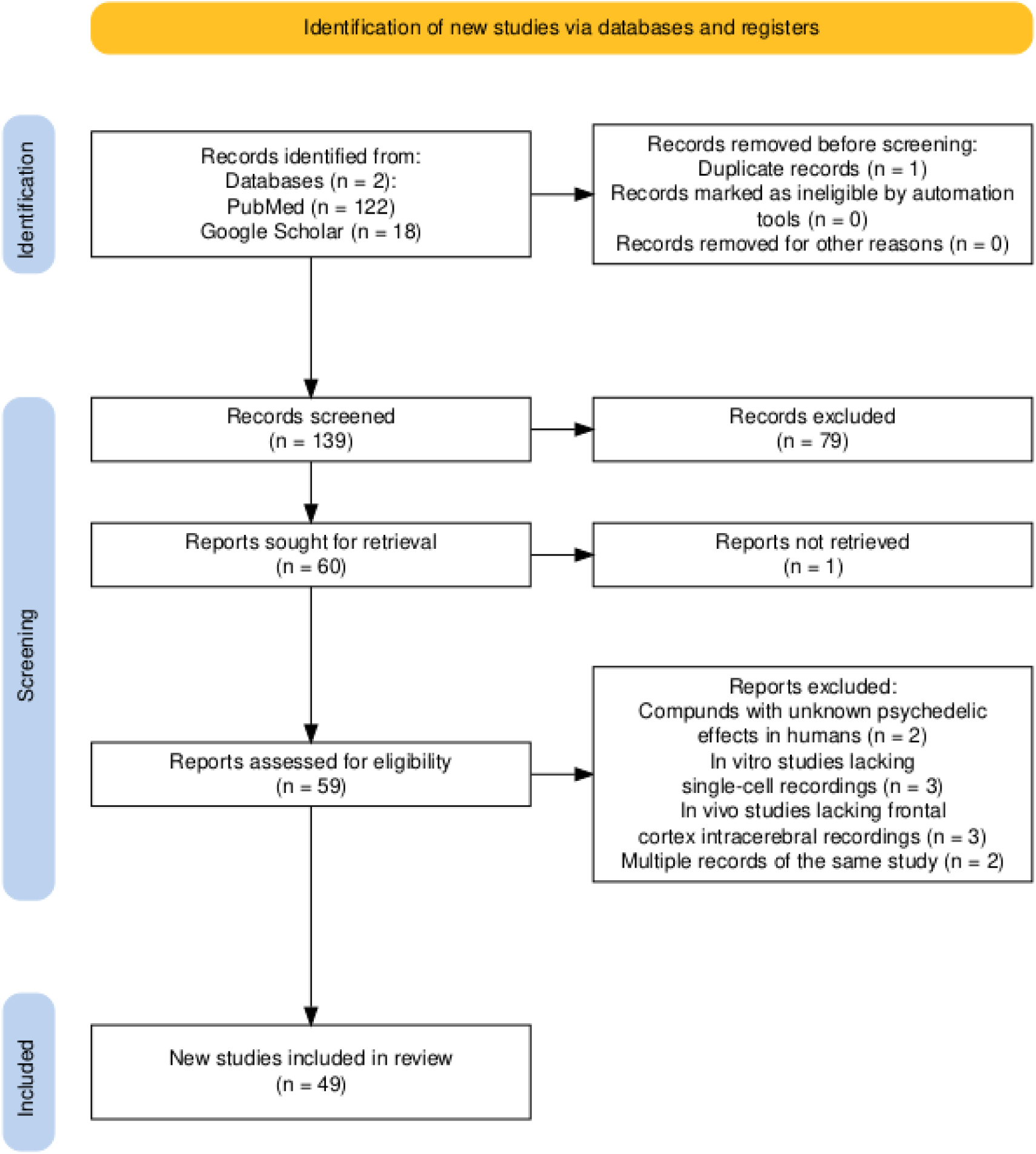
PRISMA flow diagram for study selection.

### 2.5 Study selection

139 references were screened based on titles and abstracts, and 79 records were excluded in screening, leaving 60 potential research papers. Full-text reports of the 60 identified research papers were assessed according to eligibility criteria. The systematic search identified 49 eligible studies in total: 23 *in vitro* studies, and 26 *in vivo* studies. When multiple records referred to the same underlying study (e.g., thesis, preprint, and published article), only the most complete or peer-reviewed version was retained. These duplicates were removed during the deduplication stage or consolidated at the data extraction stage, as appropriate.

## 3. RESULTS

### 3.1 In Vitro Studies

Results of *in vitro* studies are compiled in Table 1.

**Table 1:**
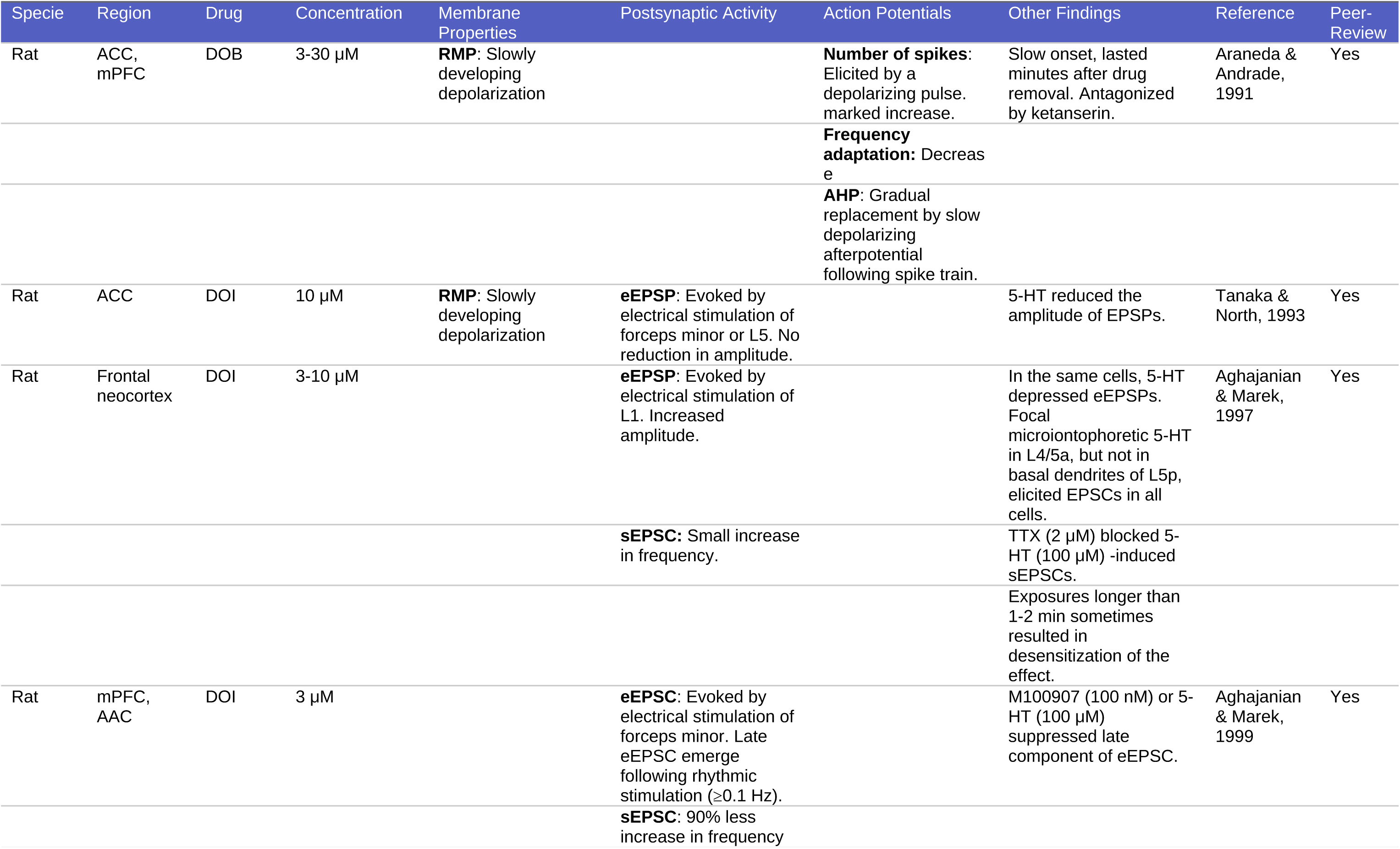
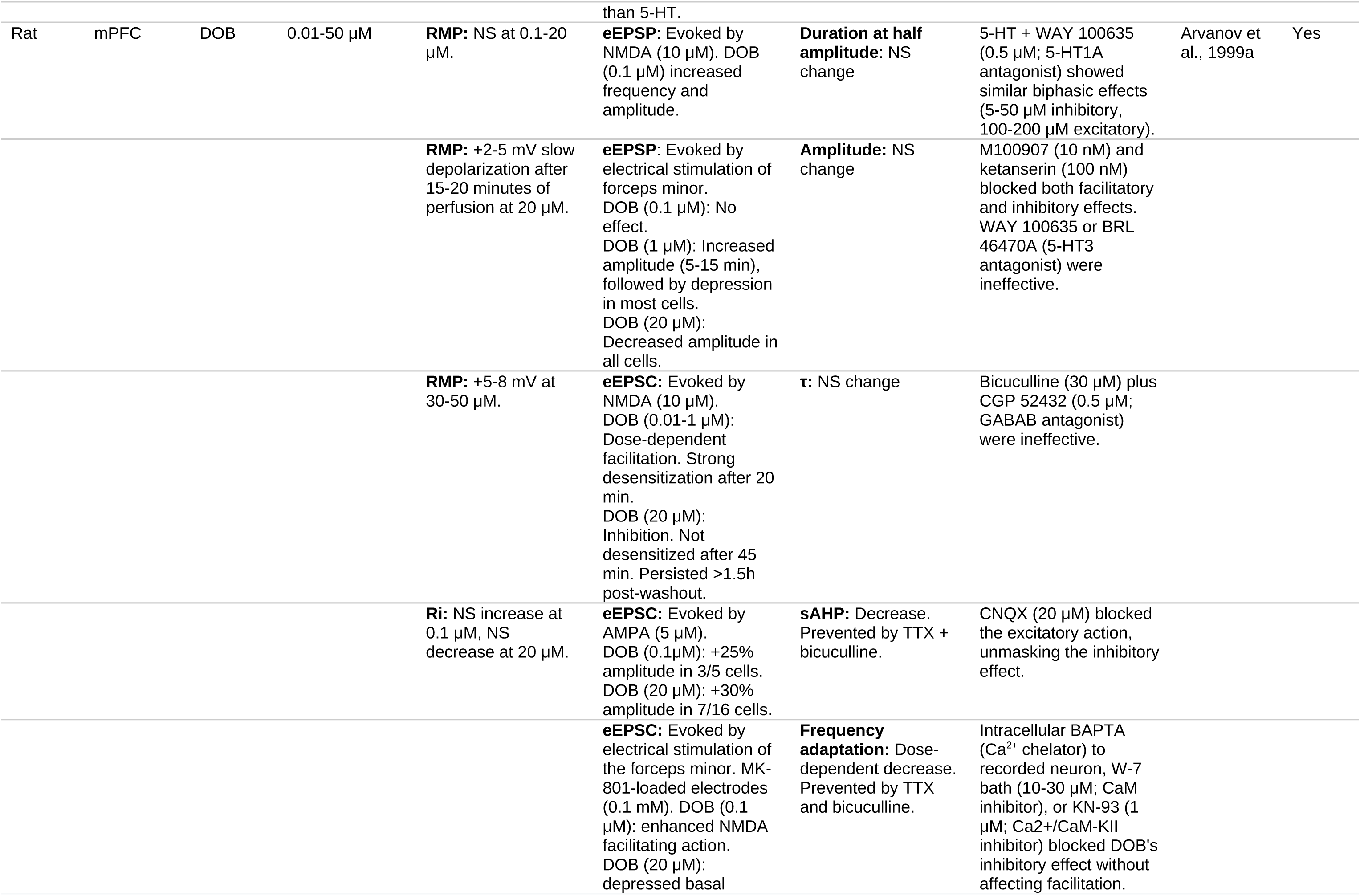
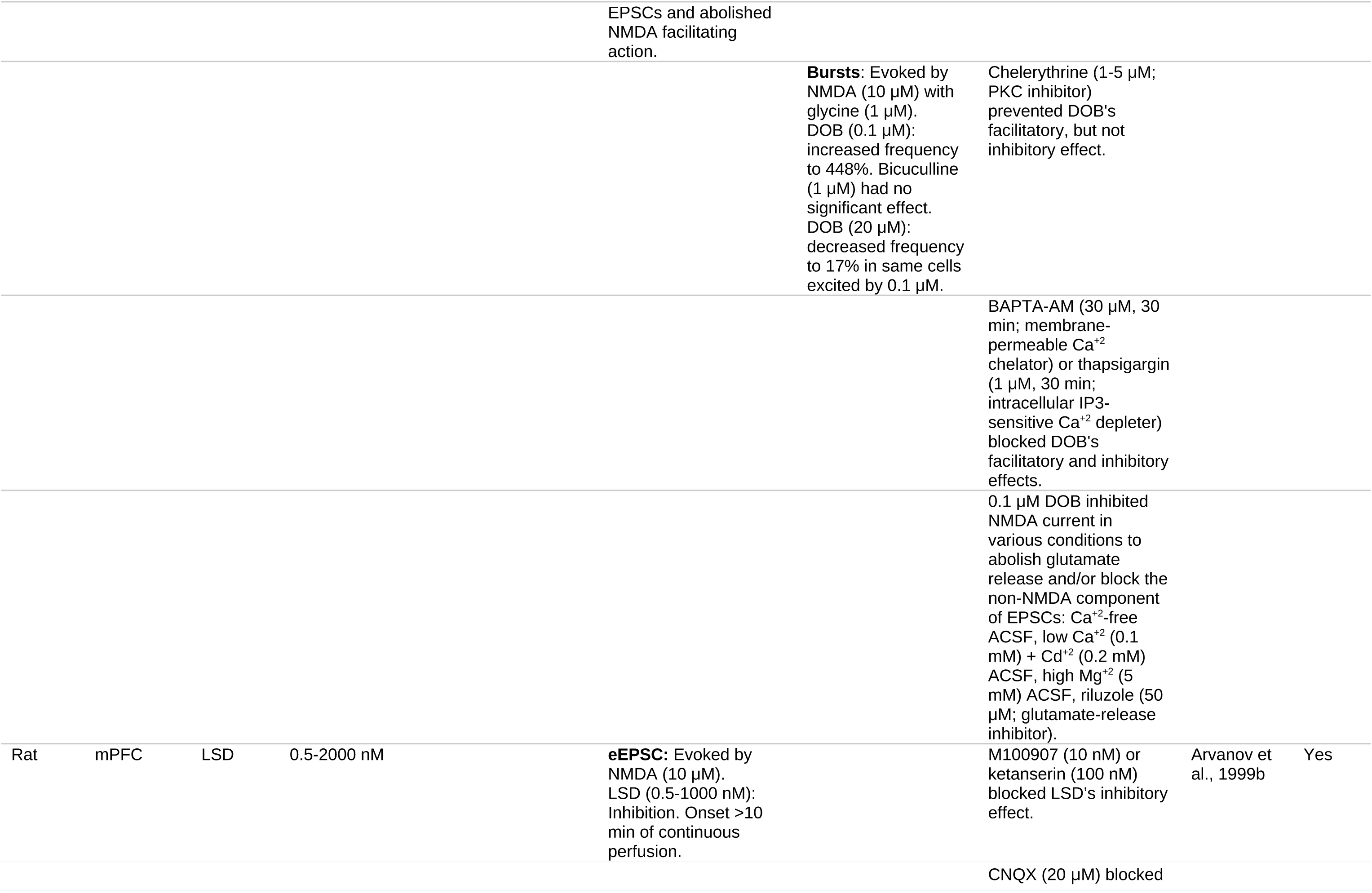
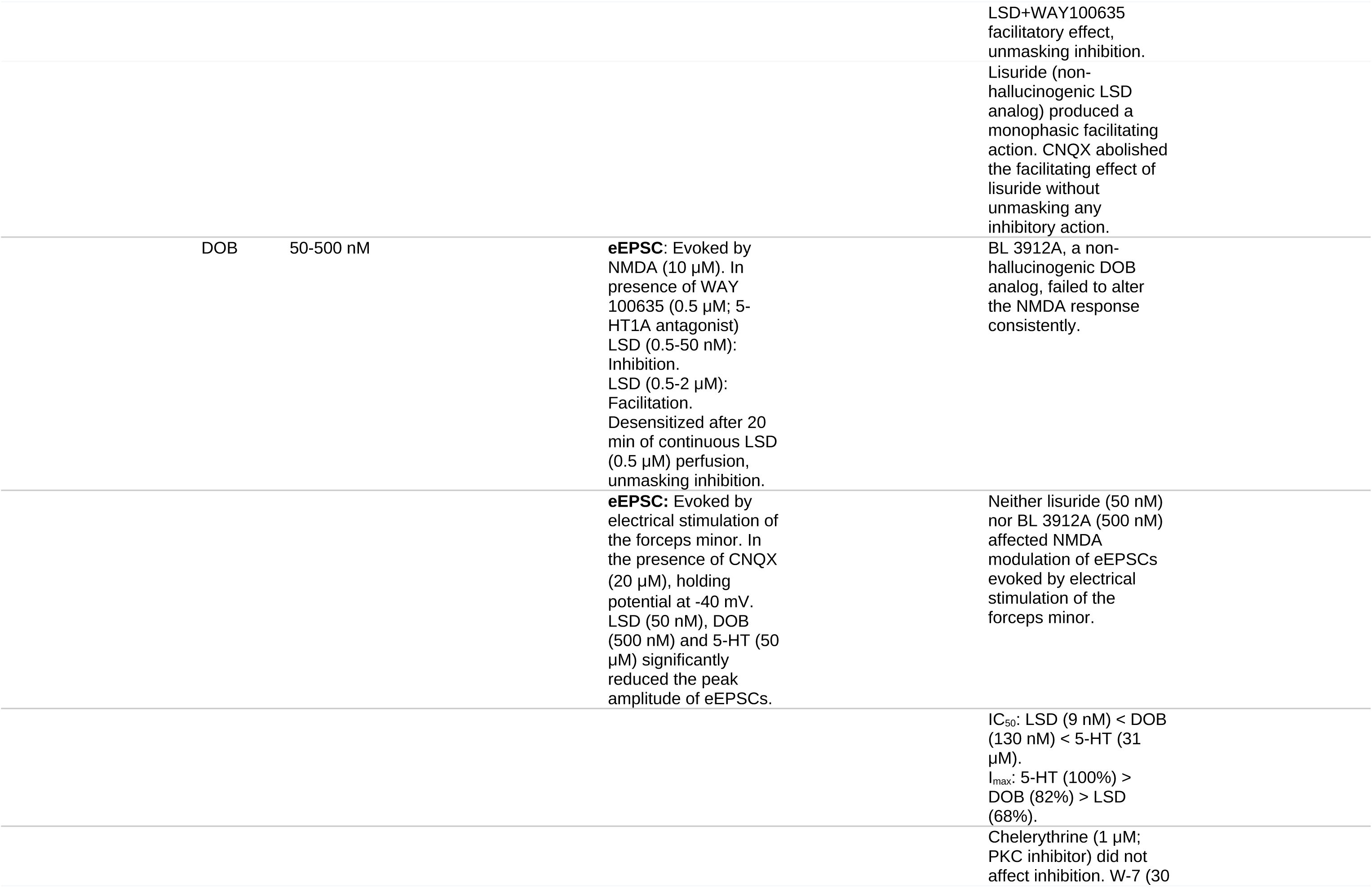
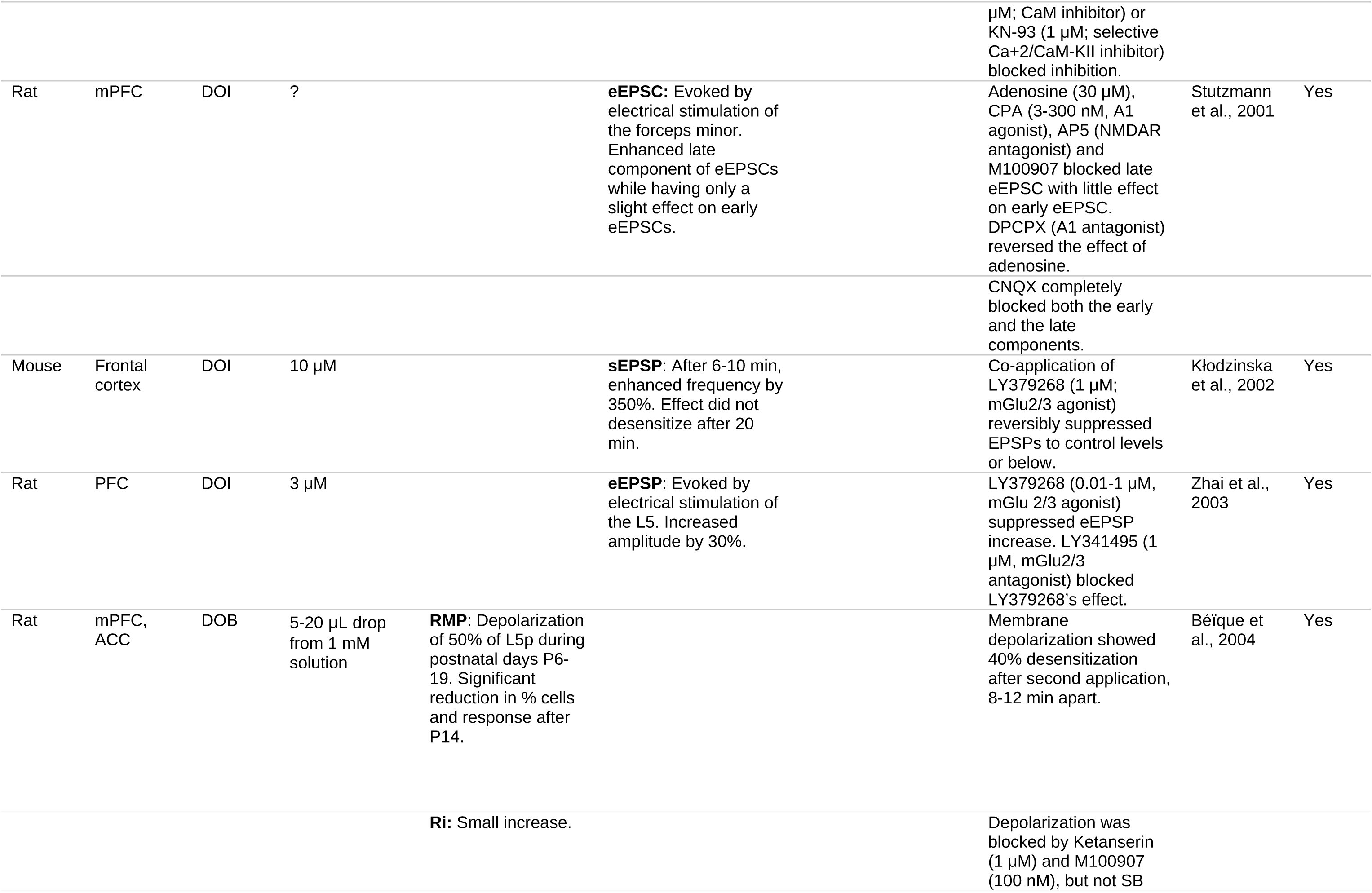
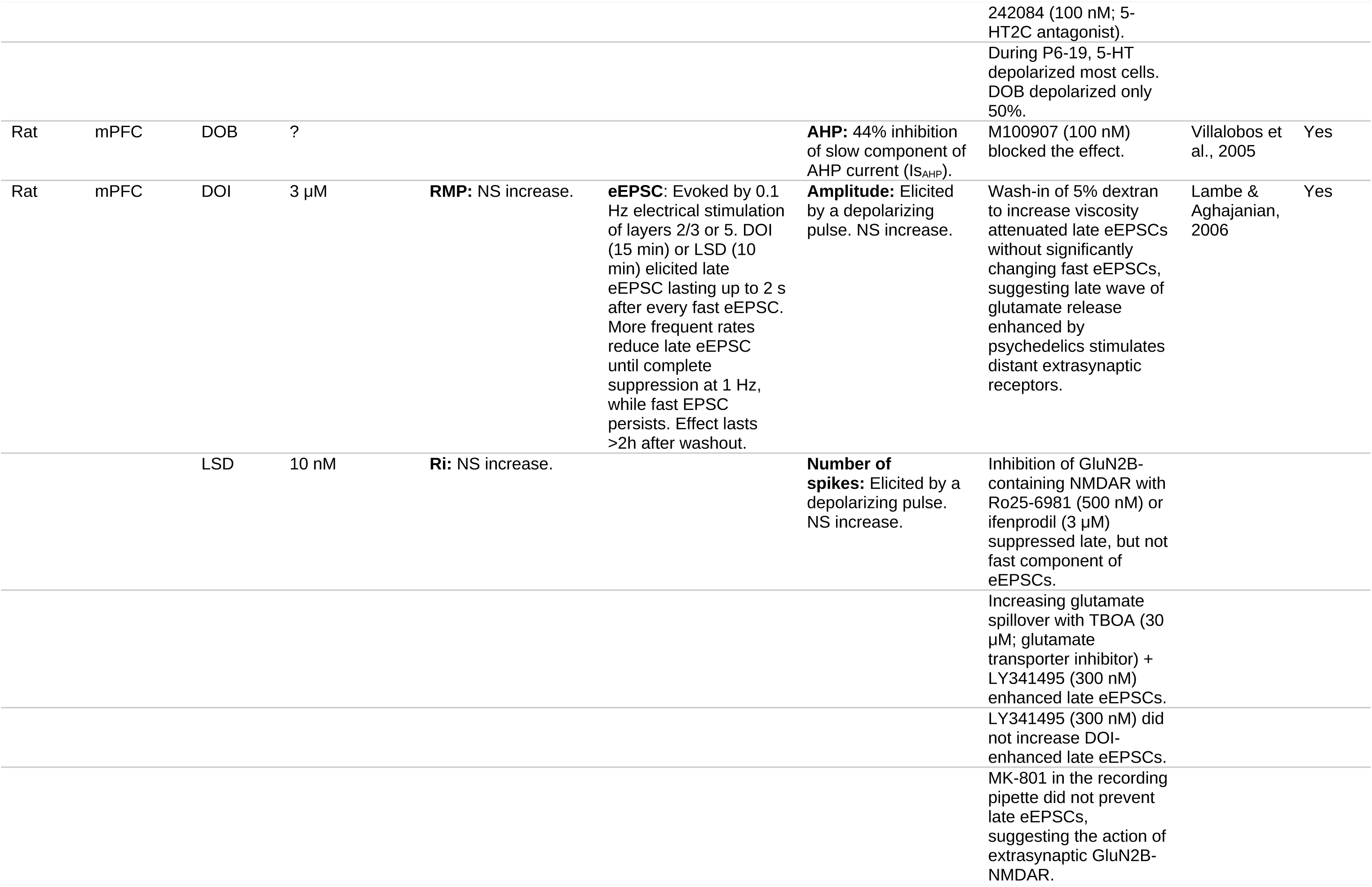
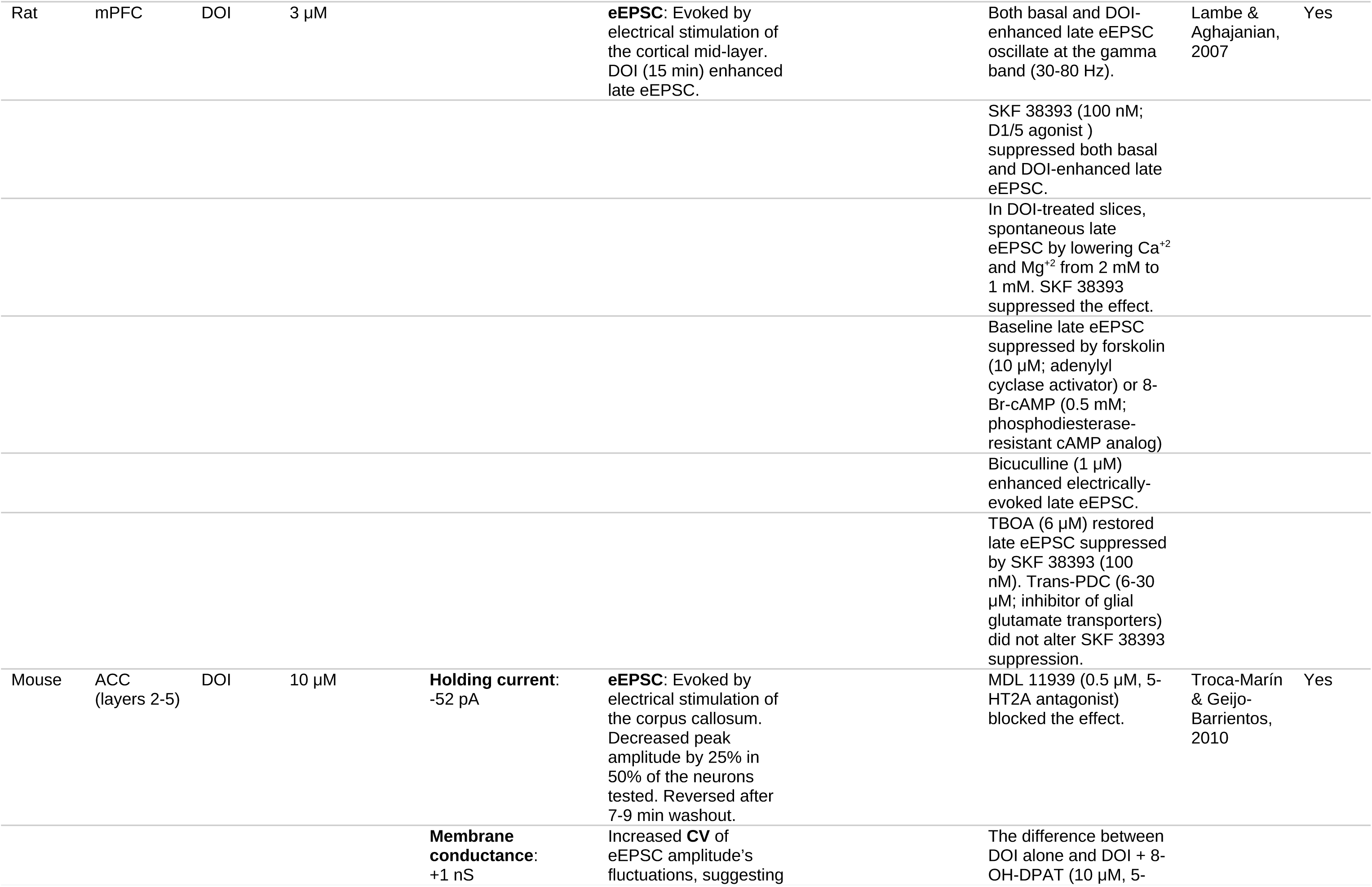
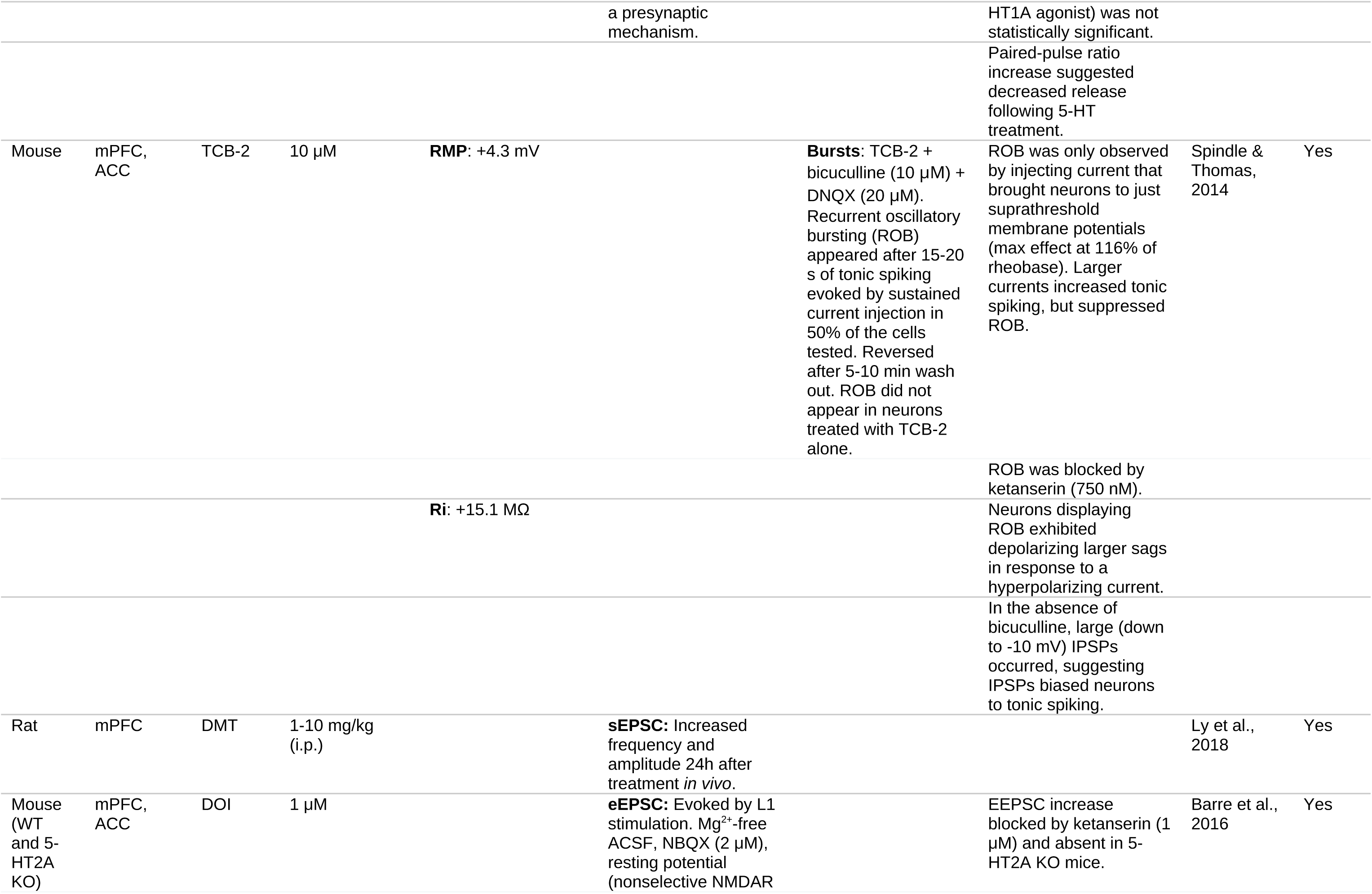
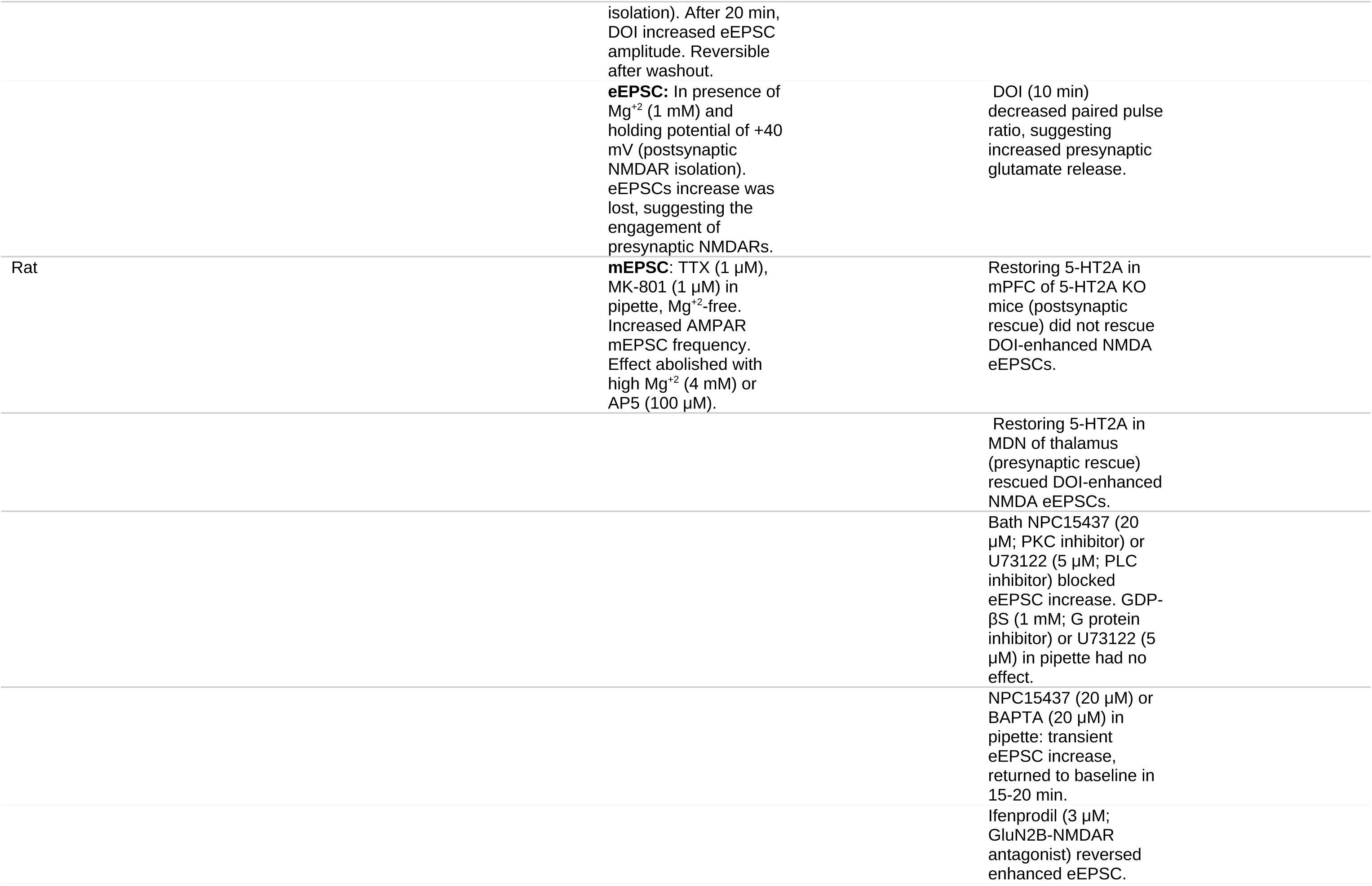
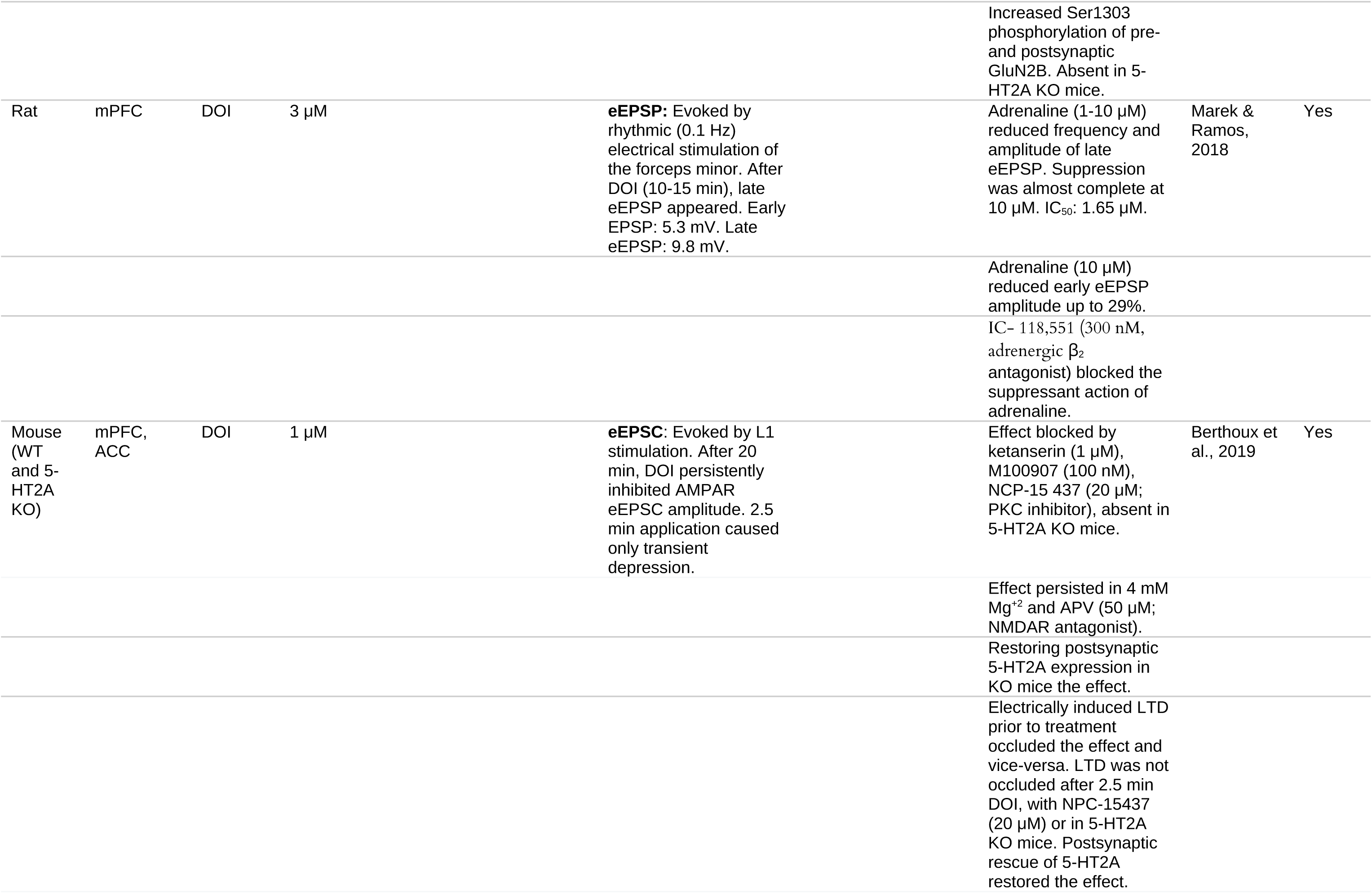
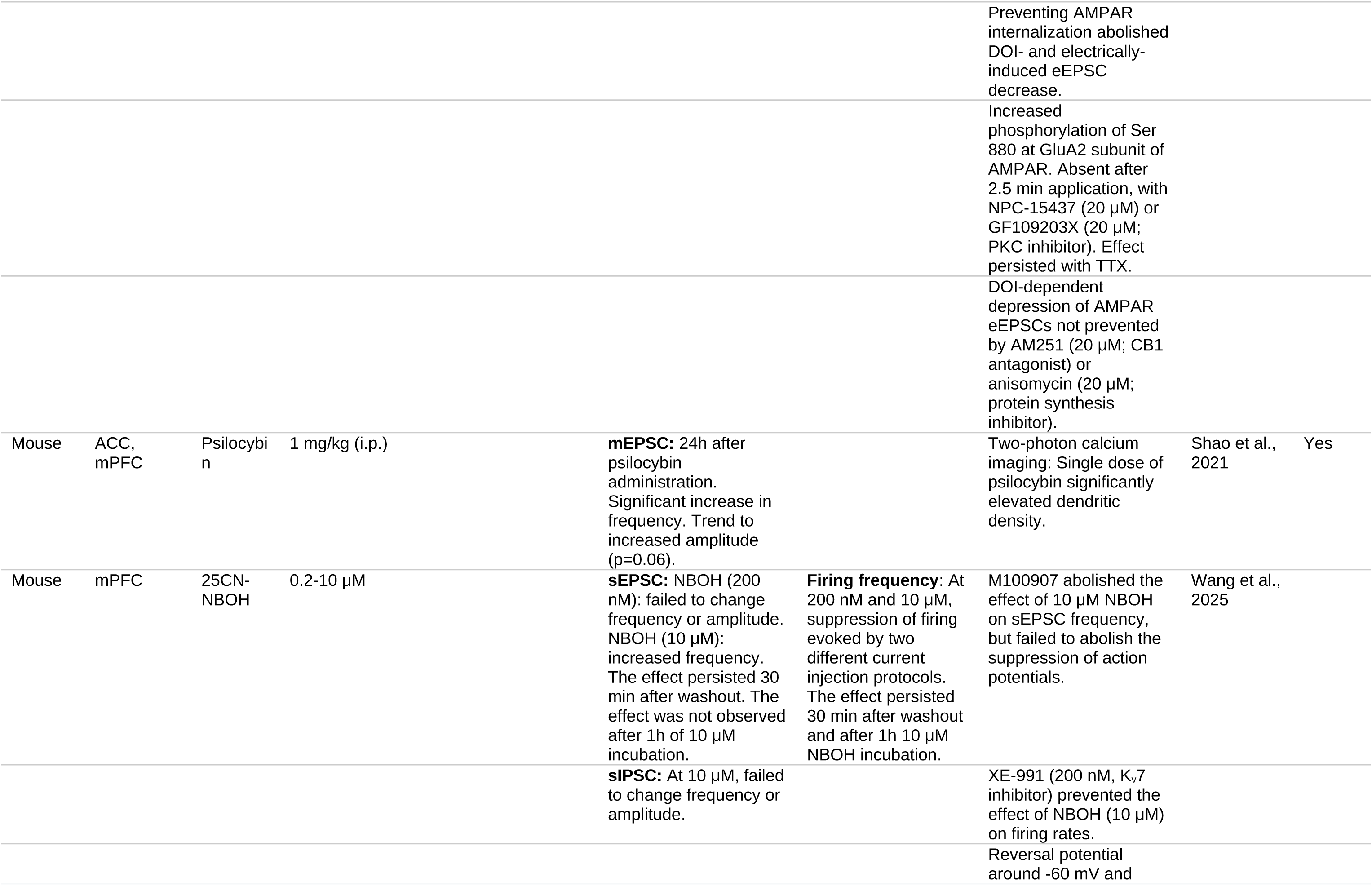
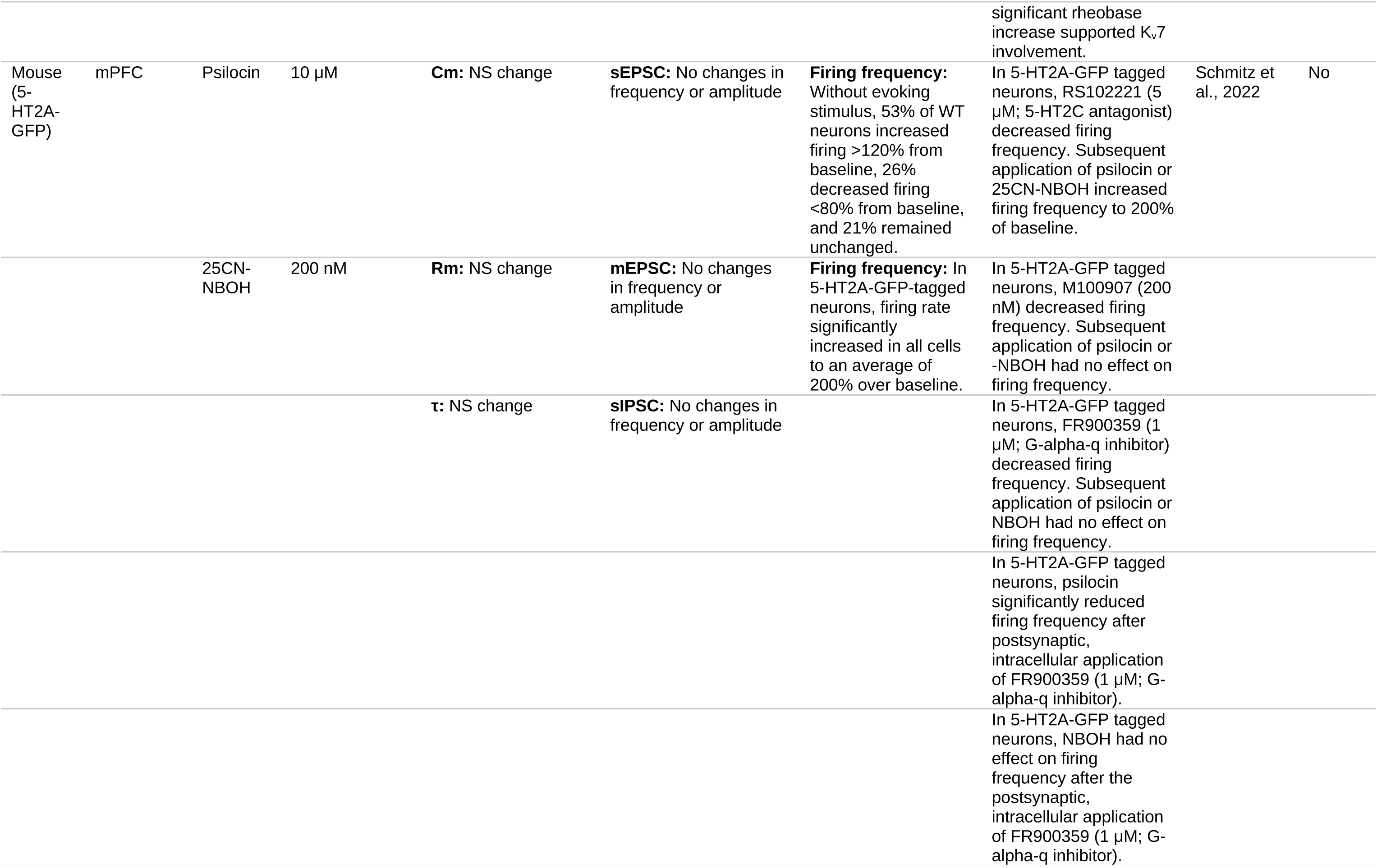
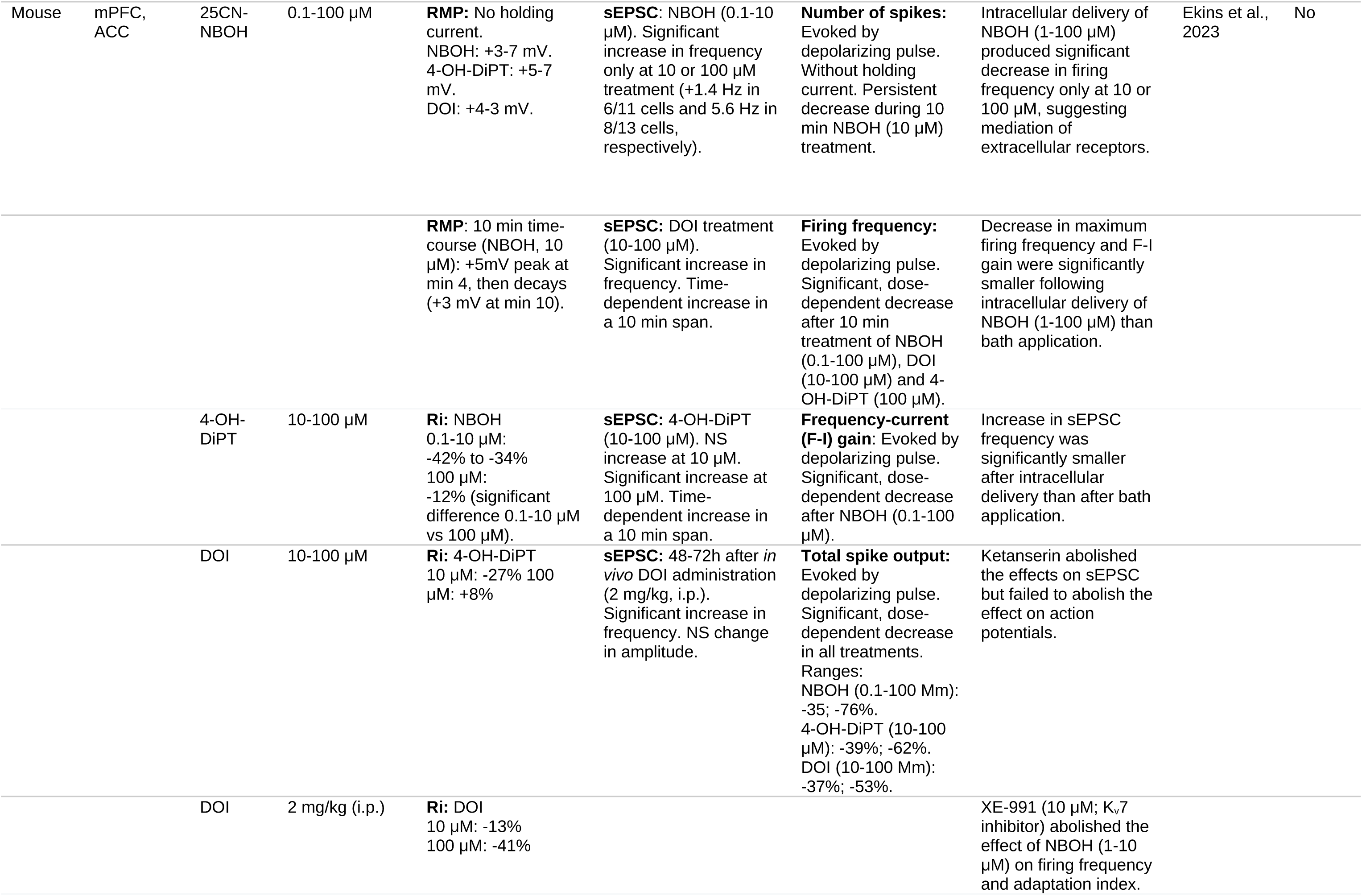
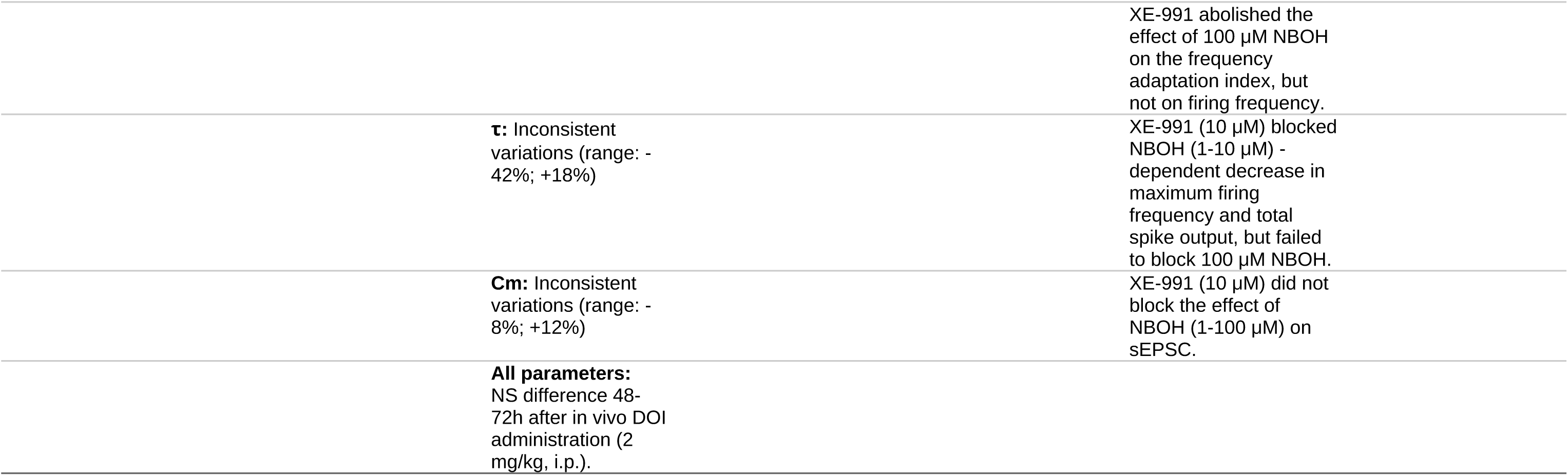
*In vitro* studies.

#### 3.1.1 Membrane Properties

##### 3.1.1.1 Resting Membrane Potential

DOB and DOI have been repeatedly shown to produce a slight, often statistically insignificant, depolarization of RMP. The effects are observed after 15–20 minutes of treatment and usually range from 2 to 5 mV (Araneda & Andrade, 1991; Arvanov et al., 1999a; Tanaka & North, 1993). More recent experiments have extended these results to other psychedelics, including TCB-2, NBOH and 4-OH-DiPT (Ekins et al., 2023; Lambe & Aghajanian, 2006; Spindle & Thomas, 2014). Despite being small, this effect is sustained in time, persisting after washout periods of up to 100 minutes (Lambe & Aghajanian, 2006). Moreover, the 5-HT_2A/C_ antagonist ketanserin can prevent RMP depolarization (Araneda & Andrade, 1991).

The effect on RMP depends on the L5p neuron population and the developmental stage. In rats aged 2–3 weeks, DOB significantly increased RMP in 50% of L5p neurons in rat mPFC. This effect was blocked by 5-HT_2A_ antagonists a selective 5-HT_2C_ antagonist. After postnatal day 14 (P14), there was a significant reduction in the fraction of cells responding to DOB and the amplitude of their response. Single-cell RT-qPCR in L5p neurons during P6–P15 suggests that the decrease of DOB’s depolarizing action cannot be fully explained by a reduction in 5-HT_2A_ expression, suggesting indirect mechanisms acting early in development (Béïque et al., 2004).

##### 3.1.1.2 Input Resistance

The reports about the effect of psychedelic drugs on Ri of L5p neurons are not consistent.

In the rat, DOB has a non-significant concentration-dependent biphasic effect, with Ri increasing at 0.1 μM and decreasing at 20 μM (Arvanov et al., 1999a). After a 15-minute perfusion of 3 μM DOI and a 5–100 minute washout period, Ri increased non-significantly (Lambe & Aghajanian, 2006). Additionally, non-significant DOI-dependent Ri increase has been observed in rat pups (Béïque et al., 2004).

In the mouse, Spindle and Thomas (2014) observed a significant Ri increase of 15 MΩ in mPFC slices treated with TCB-2. Conversely, Ekins et al. (2023) have reported significant Ri decreases by NBOH, 4-OH-DiPT, or DOI. The latter study observed an inverse concentration-dependent effect of NBOH: at lower concentrations (0.1–10 μM), NBOH significantly reduced Ri to a greater extent (-42% to -34%) than at 100 μM (-12%). Similarly, 10 μM 4-OH-DiPT reduced Ri by 27%, while 100 μM increased it by 8%. In contrast, 10 μM DOI reduced Ri by 13%, whereas 100 μM further reduced it by 41%. Anyway, since these drugs have binding affinities for 5-HT_2A_ in the nanomolar range, concentrations of 100 μM are well above in vivo or clinically relevant levels, risking the activation of off-target receptors (for instance, 100 μM NBOH is approximately 450,000 times its binding affinity to 5-HT_2A_) (Poulie et al., 2020).

##### 3.1.1.3 Other Parameters

In mouse mPFC slices, DOI decreased the holding current by 52 pA, and increased membrane conductance by 1 nS, in most cell recorded (Troca-Marín & Geijo-Barrientos, 2010). Membrane time constant and input capacitance were not consistently affected by NBOH, 4-OH-DiPT, or DOI in a way that reflected any pattern dependent on the drug or its concentration. Additionally, no significant changes in RMP, Ri, input capacitance, or time constant were observed in mouse mPFC slices obtained 48–72 hours after *in vivo* administration of 2 mg/kg DOI (i.p.) (Ekins et al., 2023).

One study evaluating the effects of psilocin on L5p spontaneous firing rates in mouse cortical slices found no significant differences in membrane resistance, capacitance, or time constant between neurons that were excited, inhibited, or unaffected by psilocin (Schmitz et al., 2022).

In summary, psychedelics induce slight but consistent depolarization of the RMP. This effect is sustained over time and depends on 5-HT_2A/C_ receptors. The impact of psychedelics on RMP varies with neuronal population and developmental stage, with younger rats showing stronger responses. Reports on input resistance (Ri) are inconsistent. While some studies indicate a biphasic or slight increase in Ri, others report significant decreases. Other membrane properties, such as capacitance and time constant, neither show consistent changes across studies. Additionally, *in vivo* administration of DOI does not appear to induce lasting effects on RMP, Ri, or other parameters.

#### 3.1.2 Synaptic Activity

##### 3.1.2.1 Biphasic Modulation of Evoked Excitatory Currents

DOB had a biphasic effect on the EPSCs elicited by N-methyl-D-aspartate (NMDA) in rat mPFC slices, co-treated with tetrodotoxin (TTX; a Na^+^ voltage-gated channel inhibitor that prevents action potentials), bicuculline (a GABA_A_ antagonist), and glycine (that potentiates the conductance of NMDA receptor, NMDAR) (Arvanov et al., 1999a).

At concentrations ranging from 0.01–1 μM, DOB elicited a dose and time-dependent facilitation of NMDA-evoked EPSCs (eEPSCs). Currents enhanced by 0.1 μM DOB returned to baseline after a 15-60-minute washing period. A second application of 0.1 μM DOB after the washing period was ineffective. Continuous perfusion with 0.1 μM DOB for 20 minutes inhibited NMDA currents, indicating strong desensitization to the facilitatory effect that lasted more than 1.5 hours. At 20 μM, DOB blocked NMDA eEPSCs. No desensitization to the inhibitory action of 20 μM DOB was observed after 30–45 minutes of continuous perfusion (Arvanov et al., 1999a).

Loading MK-801 (an open channel NMDAR antagonist) into the recording pipette selectively blocks postsynaptic NMDARs, while allowing the function of extrasynaptic and presynaptic NMDARs (McKay et al., 2013). Using MK-801-loaded pipettes, 0.1 μM DOB enhanced the facilitatory action of NMDA on eEPSCs evoked by the forceps minor’s stimulation. At 20 μM, DOB suppressed basal EPSCs and abolished NMDA’s facilitatory action. Further pharmacological manipulations to selectively block pre- or postsynaptic calcium signaling, IP3-dependent Ca^2+^ release, PKC, CaM-KII, glutamate release, and non-NMDAR-mediated excitatory currents revealed that the facilitatory effect was likely driven by a presynaptic mechanism. This mechanism involves PKC-mediated potentiation of presynaptic NMDARs by 5-HT_2A_, which increases presynaptic Ca^2+^ levels and facilitates glutamate release. Conversely, the inhibitory effect was attributed to the modulation of postsynaptic NMDARs by 5-HT_2A_, mediated by CaM-KII (Arvanov et al., 1999a).

Both the facilitatory and inhibitory effects on eEPSCs were blocked by 5-HT_2A_ antagonists. Blocking AMPA and kainate receptors (AMPAR/KAR) prevented the facilitatory action and unmasked an inhibitory effect of 1 μM DOB. Additionally, the eEPSCs were not affected by 5-HT_1A_ antagonism, 5-HT_3_ antagonism, GABA_B_ antagonism, or by altering glycine concentrations (Arvanov et al., 1999a).

In rat mPFC slices co-treated with TTX, glycine, bicuculline, and a GABA_B_ antagonist, LSD (0.5–1000 nM) invariably inhibited NMDA-evoked currents (Arvanov et al., 1999b). The onset of this effect required 10 minutes of continuous perfusion and was blocked by 5-HT_2A_ antagonists, or CaM-KII inhibitors, but not a PKC inhibitor. In the presence of a 5-HT_1A_ antagonist, LSD facilitated NMDA eEPSCs at high concentrations, an effect that disappeared after 20 minutes and was also blocked by 5-HT_2A_ antagonism. AMPAR/KAR antagonism abolished the facilitatory action of high doses, unmasking an inhibitory effect. Lisuride, a non-hallucinogenic analog of LSD, produced an invariant facilitatory effect under the same conditions (Arvanov et al., 1999b).

In the presence of an AMPAR/KAR antagonist and at a holding potential of -40 mV, which allows the isolation of postsynaptic NMDAR currents, LSD, DOB, and 5-HT reduced the peak amplitude of EPSCs evoked by electrical stimulation of the forceps minor (Arvanov et al., 1999b). The psychoactive potency of LSD, DOB, and 5-HT correlated with their IC_50_ values for inhibiting NMDAR currents: LSD (9 nM) > DOB (130 nM) > 5-HT (31 μM). Moreover, LSD’s IC_50_ aligns with its peak plasma levels in humans. Therefore, the researchers hypothesized that the inhibitory action on NMDAR currents is linked to the hallucinogenic effects of psychedelics (Arvanov et al., 1999b).

Notably, one study found that 10 μM DOI decreased the peak amplitude of eEPSCs elicited by electrical stimulation of the corpus callosum in half of the neurons tested (Troca-Marín & Geijo-Barrientos, 2010). The effect was prevented by selective 5-HT_2A_ receptor antagonism. Moreover, it was reversed after 7–9 minutes of washout, and both the coefficient of variation of amplitude fluctuations and paired-pulse ratios suggested the involvement of a presynaptic mechanism (Troca-Marín & Geijo-Barrientos, 2010). Therefore, some fibers may also elicit inhibitory actions during the early phase of psychedelic modulation. Nevertheless, L5p neurons were not isolated from other layer neurons in this assay.

##### 3.1.2.2 Mechanisms of EPSC Modulation

When NMDAR currents were non-selectively isolated in mouse mPFC slices, DOI (1 μM, 20 min) significantly increased the amplitude of EPSCs evoked by the electrical stimulation of L1 (Barre et al., 2016). This effect was rapidly reversible upon washout, absent in 5-HT_2A_ knockout (KO) mice, and blocked by ketanserin or ifenprodil, a selective antagonist of NMDARs containing GluN2B subunit. By contrast, when postsynaptic NMDAR currents were selectively isolated, DOI’s facilitatory effect was lost. Moreover, DOI decreased the paired-pulse ratio, suggesting increased presynaptic glutamate release, and, in 5-HT_2A_ KO mice, selectively restoring 5-HT_2A_ expression in thalamic nuclei targeting L5p neurons rescues the facilitatory action (Barre et al., 2016). These results support the hypothesis of psychedelics facilitating NMDAR currents through a presynaptic mechanism (Arvanov et al., 1999a).

In further agreement with Arvanov et al. (1999a), the presynaptic mechanism involved the activation of PKC. Non-selective application of PKC and PLC inhibitors blocked DOI’s facilitation, whereas postsynaptic delivery of G-protein inhibitor or PLC inhibitors had no effect. The potentiation was associated with the phosphorylation of Ser1303 in GluN2B subunits of NMDAR, which increased the receptor’s conductance and was absent in 5-HT_2A_ KO mice (Barre et al., 2016). However, unlike Arvanov et al. (1999a) findings, DOI’s facilitatory action did not desensitize after 20 minutes as long as postsynaptic Ca^2+^ levels remained high and postsynaptic PKC was not inhibited (Barre et al., 2016).

In a follow-up study, researchers isolated AMPAR currents by increasing Mg^2+^ levels and holding L5p neurons at -60 mV. DOI (1 μM, 20 min) decreased the amplitude of AMPAR EPSCs evoked by L1 electrical stimulation (Berthoux et al., 2019). The effect lasted more than 50 minutes. Shorter treatments (2.5 min) produced a transient effect that dissipated after a few minutes. AMPAR inhibition was blocked by 5-HT_2A_ antagonists or a PKC inhibitor, was unaffected by an NMDAR antagonist, and absent in 5-HT_2A_ KO mice. Restoring postsynaptic 5-HT_2A_ expression in L5p neurons rescued the effect. DOI’s action was abolished by postsynaptic PKC inhibition (Berthoux et al., 2019). Again, these results align with some of the observations of Arvanov et al. (1999a): the inhibition of eEPSCs involved a slow-onset, long-lasting, and postsynaptic mechanism. However, there are some differences, too. The inhibition of AMPAR currents involved postsynaptic PKC activity, whereas the inhibition of NMDAR currents depended on postsynaptic CaM-KII.

The modulation of AMPAR currents involved the receptor’s internalization. Electrically induced long-term depression (LTD), that is mediated by postsynaptic AMPAR internalization, occluded DOI’s effect and vice versa (Berthoux et al., 2019). Introducing an interfering peptide to block the interaction between the AMPAR subunit GluA2 and the AP2 clathrin adaptor complex, thereby preventing AMPAR internalization, abolished both LTD and DOI’s inhibitory effect. The internalization was associated with the phosphorylation of Ser880 on GluA2 (Berthoux et al., 2019).

##### 3.1.2.3 Late eEPSCs

Application of DOI (3 μM, 15 min) or LSD (10 nM, 10 min) to rat mPFC slices induced a late, prolonged (up to 2 s) component in eEPSCs (Aghajanian & Marek, 1999; Lambe & Aghajanian, 2006, 2007; Stutzmann et al., 2001). This late eEPSC is observed upon electrical stimulation of the forceps minor or the cortical midlayer, at intervals of 10 seconds or more (<0.1 Hz). It is reduced at shorter stimulation intervals and is completely suppressed at intervals of 1 second (1 Hz) (Aghajanian & Marek, 1999; Lambe & Aghajanian, 2006). Late eEPSCs remains observable for more than 2 hours after drug washout. In the absence of psychedelics, the late eEPSC occurs only occasionally, appearing in the first 10 sweeps, it has a shorter duration and is more likely to manifest at intervals of 20–30 seconds (Lambe & Aghajanian, 2006).

This mode, in which dendrites generate a prolonged depolarization after eEPSCs, corresponds to up states, which are driven by network activity (Lambe & Aghajanian, 2007). Up and down states are dendritic slow oscillations (∼1 Hz) between two membrane potentials that are determined by the precise balance of excitation and inhibition (E/I) (Larkum, 2022). When a neuron enters the up state (naturally or after psychedelics), the late eEPSC is not smooth but instead contains gamma-band oscillations (30–80 Hz) nested within the larger eEPSC waveform (Lambe & Aghajanian, 2007).

Interestingly, 5-HT does not induce late eEPSCs and, at 100 μM, it suppressed the late component elicited by DOI. The late eEPSC was suppressed by 5-HT_2A_, non-selective NMDAR and GluN2B-NMDAR antagonist, and by adenosine 1 and D1/5 receptor agonists. The A1 receptor antagonist DPCPX reversed the effects of adenosine. AMPAR/KAR antagonism suppressed both the early and late components of eEPSCs. Lowering Mg^2+^ and Ca^2+^ concentrations from 2 to 1 mM, which enhanced glutamate transmission, elicited spontaneous late EPSCs in the presence of DOI (Aghajanian & Marek, 1999; Lambe & Aghajanian, 2006, 2007; Stutzmann et al., 2001).

Lambe and Aghajanian (2006) hypothesized that glutamate spillover and the asynchronous activation of extrasynaptic GluN2B-containing NMDARs underlie late EPSCs. Supporting their hypothesis, they found that increasing medium viscosity by washing in 5% dextran to hinder glutamate diffusion attenuates the late eEPSCs without affecting the early component. Blocking postsynaptic NMDAR by delivering MK-801 through the recording pipette did not prevent the late EPSC. Moreover, inhibiting the excitatory amino acid transporter, which increased glutamate spillover, combined with an mGlu II antagonist to block the autoinhibition of glutamate release, robustly enhanced the emergence of late EPSCs in the absence of DOI, despite the mGlu II antagonist did not potentiate DOI-dependent late eEPSCs.

##### 3.1.2.4 Spontaneous and Miniature Postsynaptic Currents

In rat mPFC slices, DOI increased the frequency of spontaneous EPSCs (sEPSCs) in L5p neurons. However, the increase was only 10% of that elicited by 5-HT (Aghajanian & Marek, 1997, 1999). In mouse mPFC slices, focal application of psilocin or NBOH to 5-HT_2A_-GFP-tagged neurons did not produce significant changes in sEPSCs or spontaneous inhibitory postsynaptic currents (sIPSCs) (Schmitz et al., 2022). A bath application of NBOH was neither effective in changing sIPSCs frequency nor amplitude (Wang et al., 2025).

One preprint showed that DOI or NBOH elicited significant increases in eEPSCs frequency only at high or very high concentrations (10–100 μM). Similarly, 4-OH-DiPT increased eEPSCs frequency significantly only at 100 μM. The 5-HT_2A/C_ antagonist ketanserin blocked the action of NBOH, while XE-991, a K_v_7 channel inhibitor, had no effect (Ekins et al., 2023). They found a significantly greater increase in eEPSCs frequency upon bath application of NBOH compared to intracellular drug delivery, suggesting extracellular 5-HT_2A_-mediation (Ekins et al., 2023). Similar results were reported in a recent peer-reviewed article, where NBOH failed to alter the frequency or amplitude of eEPSCs at 200 nM, but enhanced their frequency at 10 μM (Wang et al., 2025). This effect persisted 30 minutes after washout, but, notably, was lost when the slices were incubated with the drug for 1 hour. The effect was blocked by selective 5-HT_2A_ receptor antagonism (Wang et al., 2025).

However, psychedelics of the NBOMe class, including NBOH, exhibit extremely high affinity for the 5-HT_2A_ receptor. At concentrations far exceeding the nanomolar range, these compounds also act on other serotonin receptors and receptors of a different class, like adrenergic receptors (Poulie et al., 2020). Therefore, the results of Ekins et al (2023) and Wang et al. (2025) might not be relevant *in vivo*.

Miniature EPSCs (mEPSCs) were neither significantly affected by the focal application of psilocin nor NBOH (Schmitz et al., 2022). In contrast, a bath application of DOI increased the frequency of AMPAR mEPSCs (Barre et al., 2016). The effect was abolished in a high Mg^2+^ (4 mM) medium or after a bath application of an NMDAR antagonist. The increase in mEPSCs was likely due to increased presynaptic glutamate release (Barre et al., 2016).

In mPFC slices obtained from rats treated with systemic DMT 24 hours prior to sample collection, an increase in both frequency and amplitude of eEPSCs was observed (Ly et al., 2018). Similarly, in mPFC slices from mice treated with psilocybin 24 hours prior to sample collection, an increase in frequency and a trend toward increased amplitude (p = 0.06) of mEPSCs was seen (Shao et al., 2021). These late increases in eEPSCs and mEPSCs were attributed to dendritic plasticity and synaptogenesis, rather than to direct electrophysiological action (Ly et al., 2018; Shao et al., 2021).

##### 3.1.2.5 Excitatory Postsynaptic Potentials

In L5p neurons, DOI (3–10 μM) increased the amplitude of evoked EPSPs (eEPSPs) elicited by electrical stimulation of Layer 1 (L1) (Aghajanian & Marek, 1997). At 3 μM, DOI increased the amplitude of eEPSPs by 30% following L5 electrical stimulation (Zhai et al., 2003). This effect could be abolished by the metabotropic glutamate receptor II (mGlu II) agonist LY379268. In turn, the action of LY379268 was blocked by a mGlu II antagonist (Zhai et al., 2003). By contrast, at 10 μM, DOI didn’t have any significant effect on the amplitude of eEPSPs elicited by stimulation of L5p somata or the forceps minor (Tanaka & North, 1993).

In mouse cortical slices treated with picrotoxin (a GABA_A_ antagonist) to isolate excitatory input, DOI (10 μM) increased the frequency of spontaneous EPSPs (eEPSPs) in L5p cells two- to tenfold (mean 350%) after 6–10 minutes of treatment, and the effect did not desensitize over a 20-minute application. LY379268 reversed eEPSPs below control levels (Kłodzinska et al., 2002).

DOB elicited a time and concentration -dependent modulation of EPSPs that matched that of EPSCs (Arvanov et al., 1999a). At 1 μM, DOB increased eEPSP amplitude evoked by forceps minor’s stimulation in most L5/6p neurons. This facilitatory effect was lost after 5–15 minutes, followed by an inhibitory effect (Arvanov et al., 1999a). At 20 μM, DOB reduced the amplitude of eEPSPs in all cells, and this inhibitory effect is potentiated after 20 minutes, during which the RMP is slowly depolarized (Arvanov et al., 1999a).

DOI (3 μM, 10-15 min) induced the appearance of late eEPSPs following rhythmic (0.1 Hz) stimulation of forceps minor fibers (Marek & Ramos, 2018). Late eEPSPs were larger in amplitude than early eEPSPs (9.8 mV vs 5.3 mV) and were preferentially and dose-dependently suppressed by ꞵ_2_ activation (Marek & Ramos, 2018).

#### 3.1.3. Action Potentials

##### 3.1.3.1 Intrinsic excitability

The reports of psychedelic action on neural intrinsic excitability are inconsistent. Early research in rat mPFC slices reported that DOI markedly increases the number of spikes elicited by a depolarizing current pulse in L5p neurons. This effect has a slow onset and is antagonized by ketanserin (Araneda & Andrade, 1991). However, subsequent experiments found that DOI elicits only a slight, non-significant increase in the number of spikes (Lambe & Aghajanian, 2006).

In mouse mPFC slices, it has been recently shown that NBOH elicits a significant, dose-dependent decrease in total spike output from L5p neurons during a current injection (Ekins et al., 2023; Wang et al., 2025). This is accompanied by significant decreases in average firing frequency, maximum firing frequency, and firing frequency-current input gain (Ekins et al., 2023). Similar inhibitory effects were reported for DOI and 4-OH-DiPT. The loss in frequency-current gain was more pronounced when NBOH was bath-applied compared to intracellular delivery, suggesting that the action is mediated by extracellular 5-HT_2A_ (Ekins et al., 2023).

The firing frequency after combining NBOH (0.1 or 10 μM) with 5-HT_2A_ antagonists was significantly lower than with antagonists alone, suggesting that 5-HT_2A_ does not mediate the inhibitory effect at these concentrations (Ekins et al., 2023; Wang et al., 2025). In turn, XE-991 blocked the decreased firing by 1 or 10 μM NBOH, but failed to do so for 100 μM NBOH, indicating that K_v_7 channels are involved in the inhibitory action of NBOH at concentrations of 1–10 μM, but not of 100 μM (Ekins et al., 2023; Wang et al., 2025).

However, the findings of Ekins et al. (2023) and Wang et al. (2025) should be interpreted with caution. We have already noted the use of concentrations high above drug affinity for 5-HT_2A_. Additionally, pharmacological tests with 5-HT_2A_ antagonists (ketanserin or M100907) and XE-991 lack within-cell comparisons across baseline, drug, antagonist, and combined conditions, limiting causal interpretation. The absence of these controls is particularly critical given the high inter-cellular variability observed in drug responses (Ekins et al., 2023; Wang et al., 2025). Moreover, the ability of 10 μM ketanserin to effectively block 10 μM NBOH is questionable, as 2.2 nM NBOH inhibits 50% of 0.5 nM ketanserin binding to 5-HT_2A_ (Halberstadt et al., 2016).

##### 3.1.3.2 Burst Firing

DOB (0.1 μM) increased burst firing frequency elicited by NMDA by 448%. Bicuculline does not prevent the increase, suggesting that it is not due to interneuron disinhibition. By contrast, 20 μM DOB decreased bursting frequency to 17% in the same cells that were excited by 0.1 μM DOB (Arvanov et al., 1999a).

In mouse mPFC slices treated with GABA_A_ and AMPAR/KAR antagonists to isolate L5p neurons from both inhibitory and excitatory inputs, TCB-2 elicited recurrent oscillatory bursts (ROBs) after 15–20 s of tonic spiking induced by a depolarizing current, in 50% of the cells tested. Neurons that exhibited ROBs had significantly larger depolarizing sags in response to hyperpolarizing currents. ROBs were observed when neurons are held at just suprathreshold potentials, with the maximum effect at 116% of rheobase. They were blocked by ketanserin and disappeared after 5–10 min of washout (Spindle & Thomas, 2014).

ROBs did not appear in the absence of GABAR and AMPAR/KAR antagonists. In the absence of GABA_A_ antagonists, large inhibitory postsynaptic potentials (IPSPs), down to - 10 mV, appeared to bias neurons toward tonic spiking when entering the ROB mode. Similar ROB discharges have been reported following the focal application of glutamate to the apical dendrites of L5p or could be elicited by concurrent stimulation of the apical dendrites and soma of L5p (Spindle & Thomas, 2014).

##### 3.1.3.3 Spontaneous Firing Frequency

Focal application of psilocin to untagged L5p neurons within cortical slices increased the firing frequency in 53% of cells, while 26% were inhibited and the rest remained unchanged. In this experiment, excitation and inhibition were defined as ±20% changes from the baseline firing frequency. By contrast, in L5p neurons tagged with a 5-HT_2A_-GFP fusion protein, psilocin or NBOH increased the firing frequency by 200% in all cells (Schmitz et al., 2022).

When 5-HT2C receptors were antagonized, the basal firing frequency decreased. Subsequent application of psilocin or NBOH reversed the decrease and raised the firing frequency by 200%. Focal application of either a 5-HT2A antagonist or a Gαq/11 inhibitor also reduced basal firing frequency, but this decrease could be reversed by psilocin or NBOH (Schmitz et al., 2022).

The intracellular delivery of the Gα_q/11_ inhibitor made psilocin reduce the firing frequency of L5p neurons, which was not observed with NBOH. The discrepancy can be explained by the fact that psilocin can still recruit Gα_i/o_ signaling via 5-HT_1A_ receptors in the presence of Gα_q/11_ inhibitors, while NBOH is selective for Gα_q/11_-coupled 5-HT receptors (Schmitz et al., 2022).

##### 3.1.3.4 Spike Frequency Adaptation, Afterhyperpolarization and Spike Amplitude

Several studies report that DOB decreases spike frequency adaptation in trains of action potentials elicited by a depolarizing pulse in rat mPFC slices (Araneda & Andrade, 1991; Arvanov et al., 1999a). This effect could be prevented by TTX plus bicuculline (Arvanov et al., 1999a).

The findings from the preprint by Ekins et al. (2023) conflict with these results, showing a significant, concentration-dependent increase in frequency adaptation after NBOH treatment in mouse mPFC slices. The adaptation index was not significantly different after combining XE-991 with NBOH, compared to XE-991 alone, suggesting that blocking K_v_7 channels occludes frequency adaptation. However, these pharmacological manipulations lack a comparison with NBOH treatment alone and vehicle controls.

Spike afterhyperpolarization (AHP) is mediated by the opening of K^+^ channels and is closely linked to frequency adaptation. DOI or DOB decreased AHP in L5p neurons from rat mPFC slices (Araneda and Andrade, 1991; Arvanov et al., 1999a; Villalobos et al., 2005), with one study reporting the replacement of AHP by a slow depolarizing afterpotential following a train of spikes (Araneda & Andrade, 1991). AHP can be decomposed in an early and a slowly-decaying component. DOB reduced the latter along with the current that mediates it, an action that could be blocked by 5-HT_2A_ antagonism (Arvanov et al., 1999a; Villalobos et al., 2005).

Spike amplitude modulation has been inconsistently reported. Some studies in rat cortical slices report non-significant increases or decreases after DOI or DOB treatments (Arvanov et al., 1999a; Lambe & Aghajanian, 2006). In mouse cortical slices, Ekins et al. (2023) report a significant, concentration-dependent decrease in spike amplitude (-2% to -19%) following 0.1–100 μM NBOH. Similarly, 4-OH-DiPT at 10 μM caused a 2% decrease in spike amplitude, while 100 μM caused a 12% decrease. In turn, DOI showed an 8% decrease in spike amplitude at 10 μM but only a 4% decrease at 100 μM.

### 3.2 In Vivo Studies

Results of *in vivo* studies are compiled in Table 2.

**Table 2:**
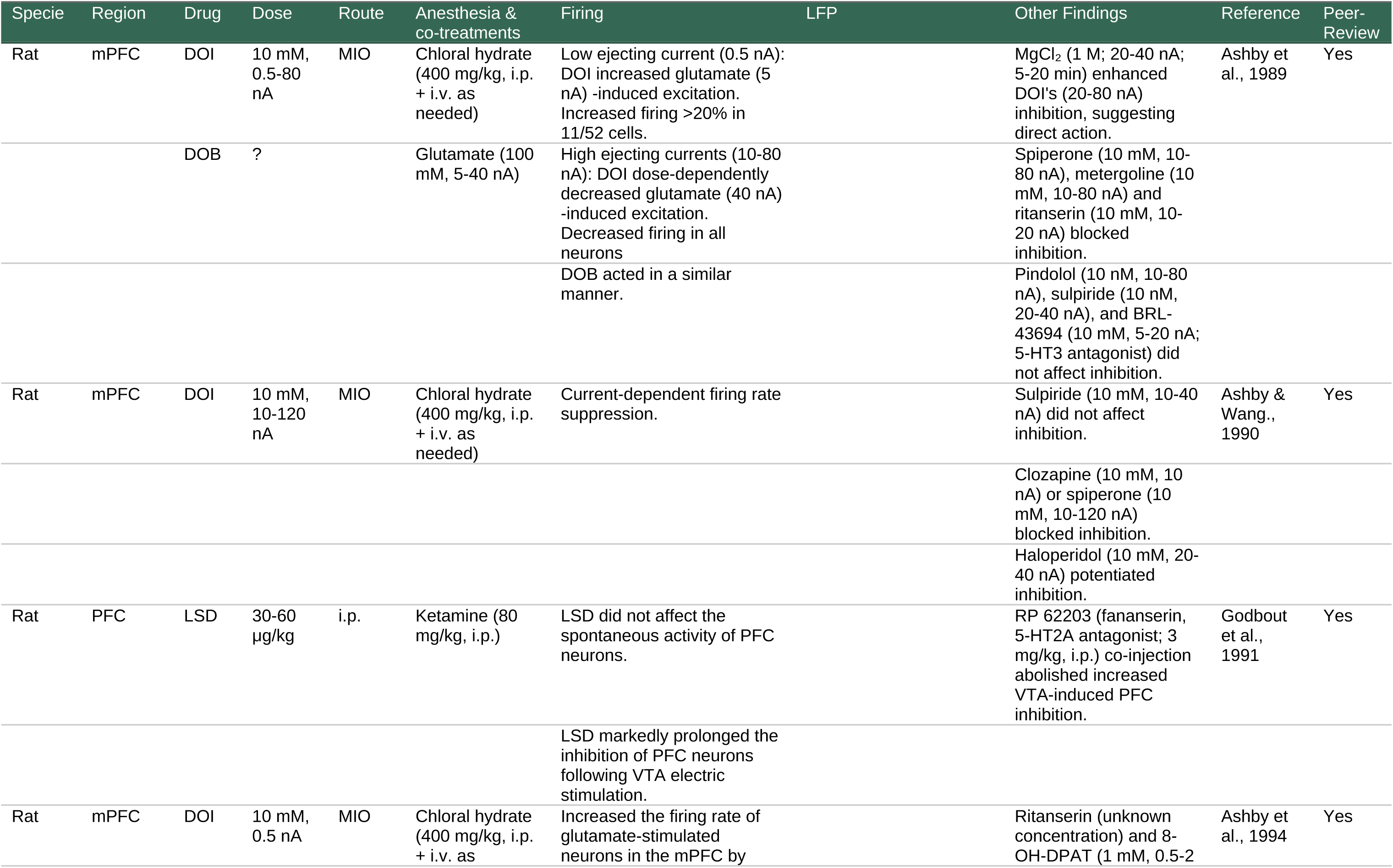
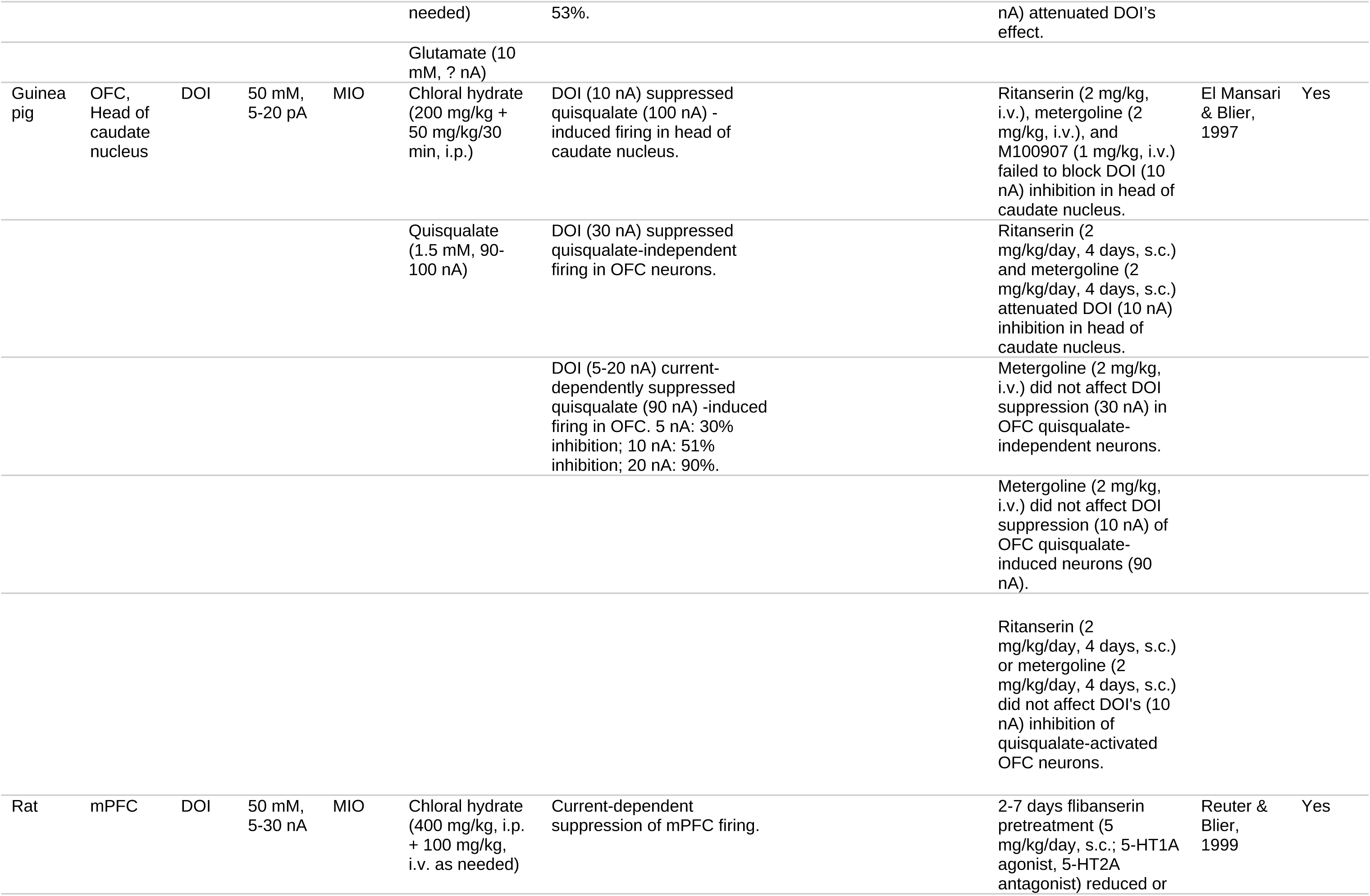
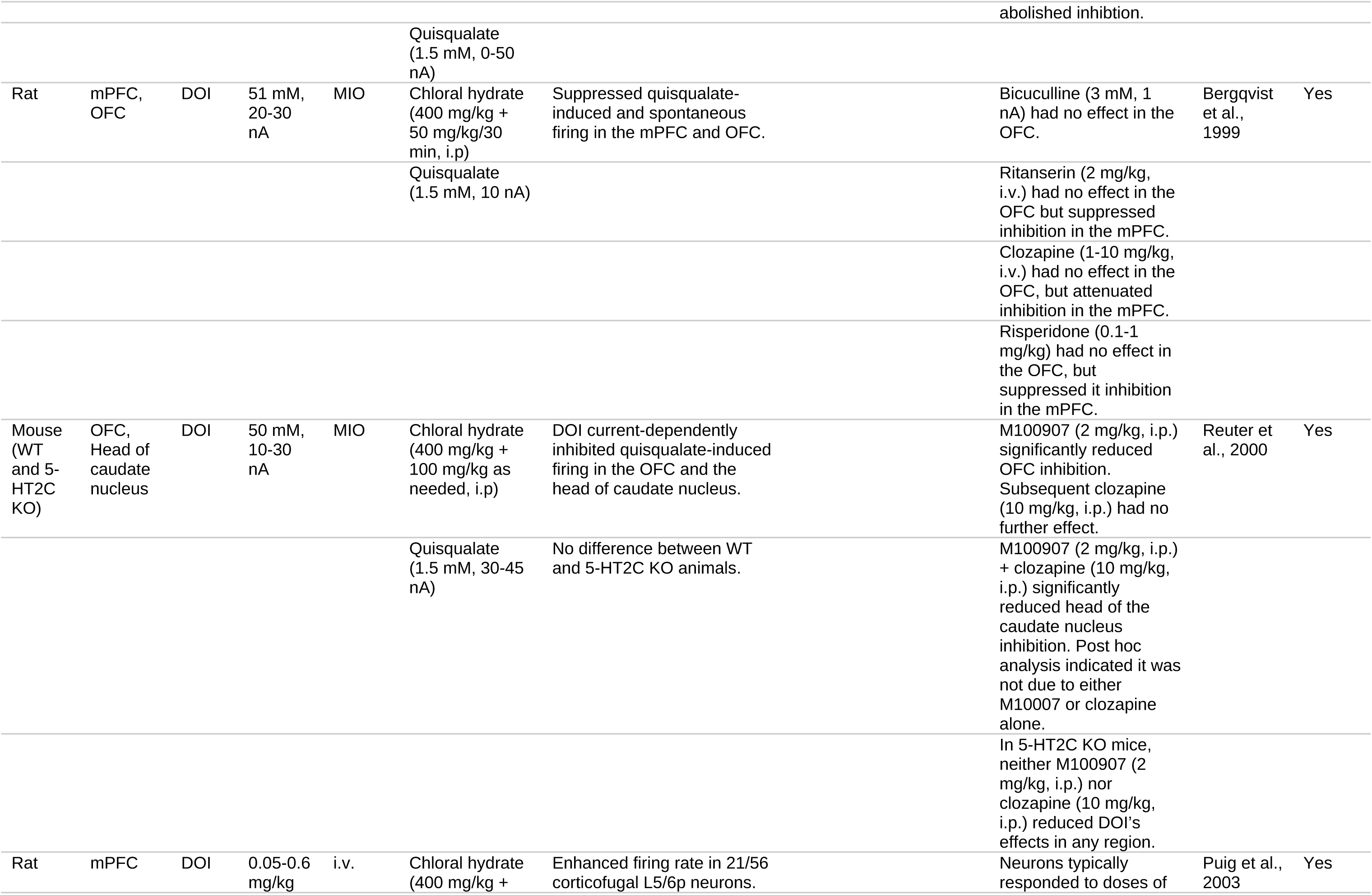
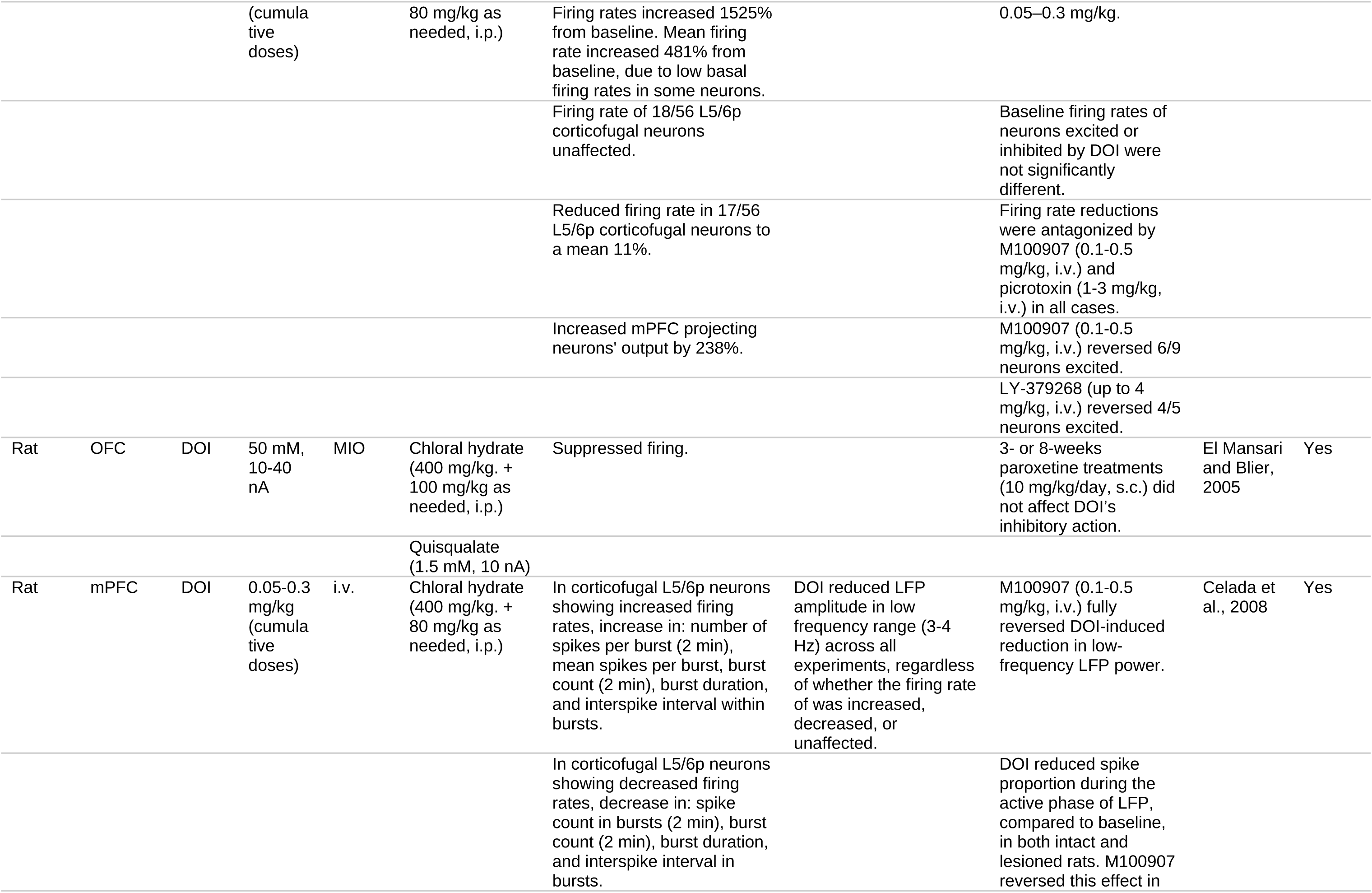
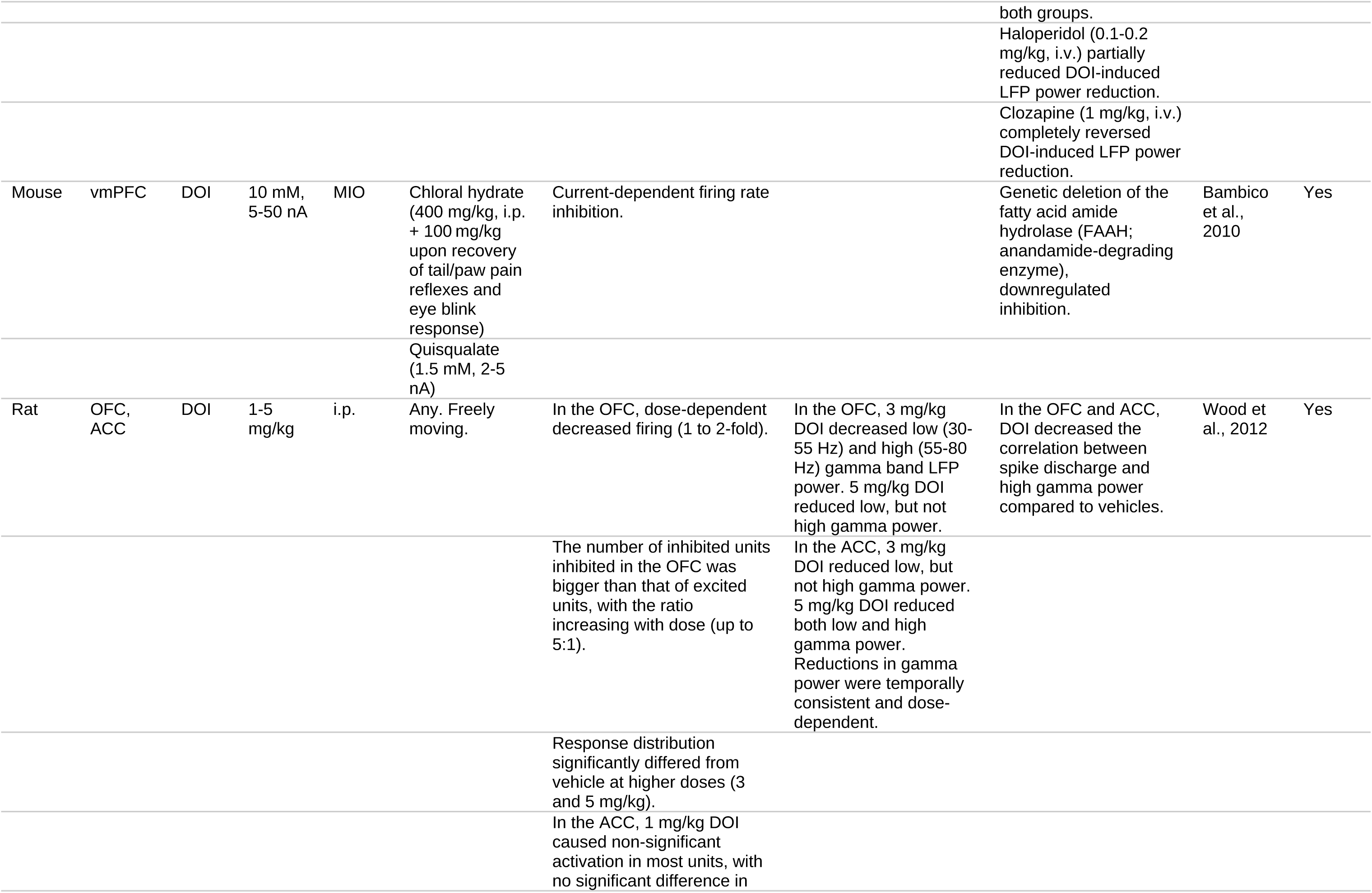
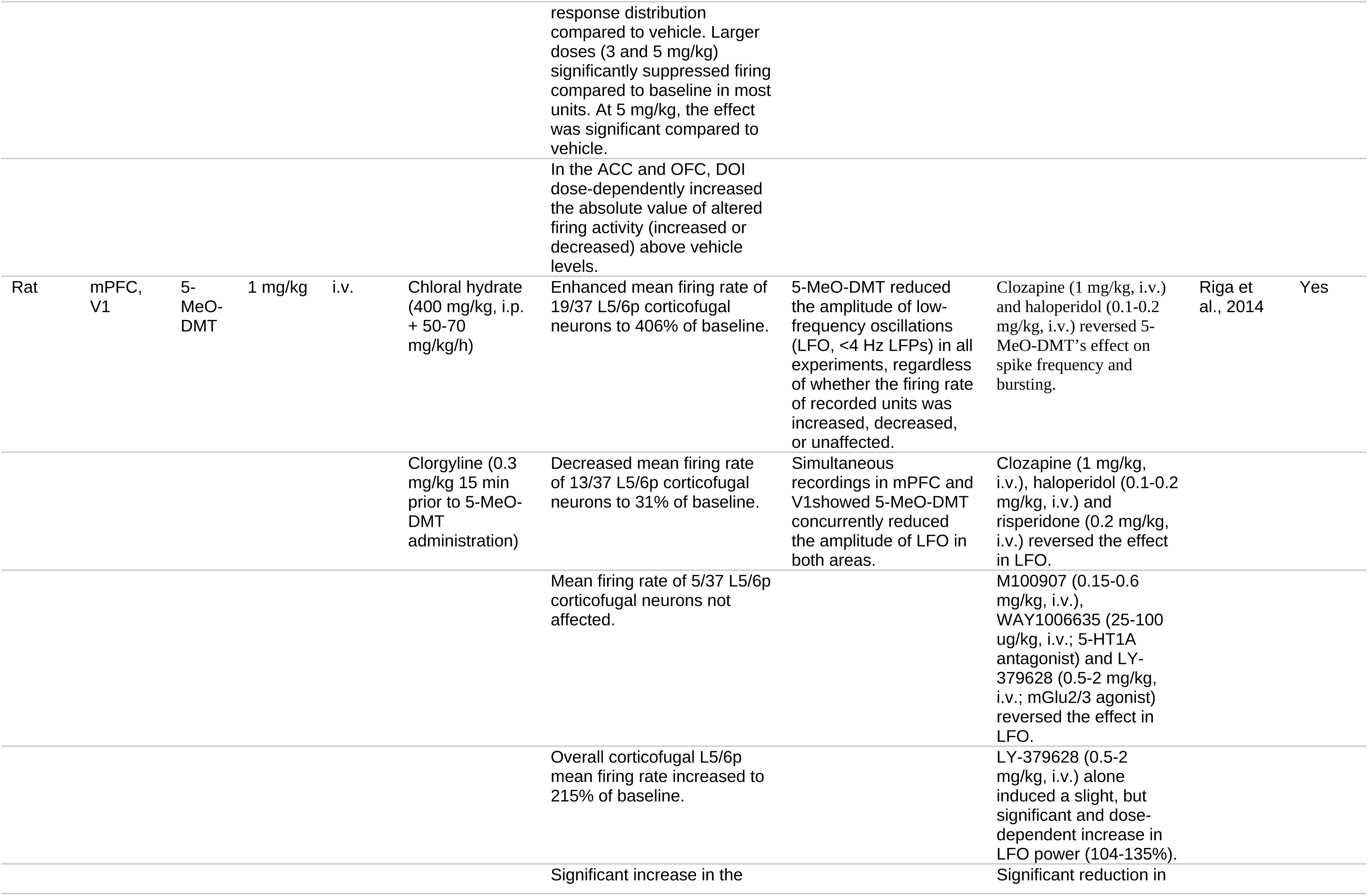
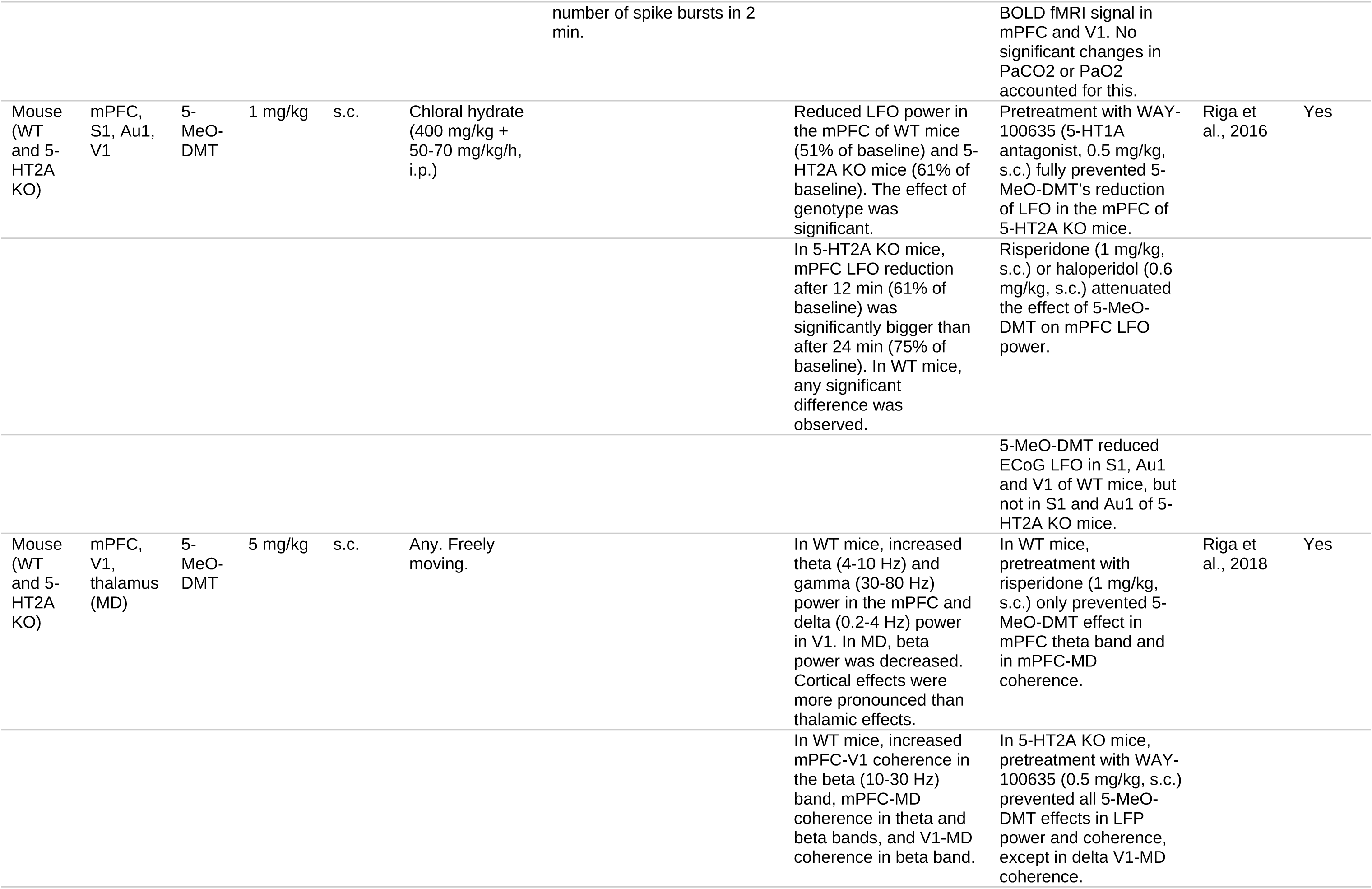
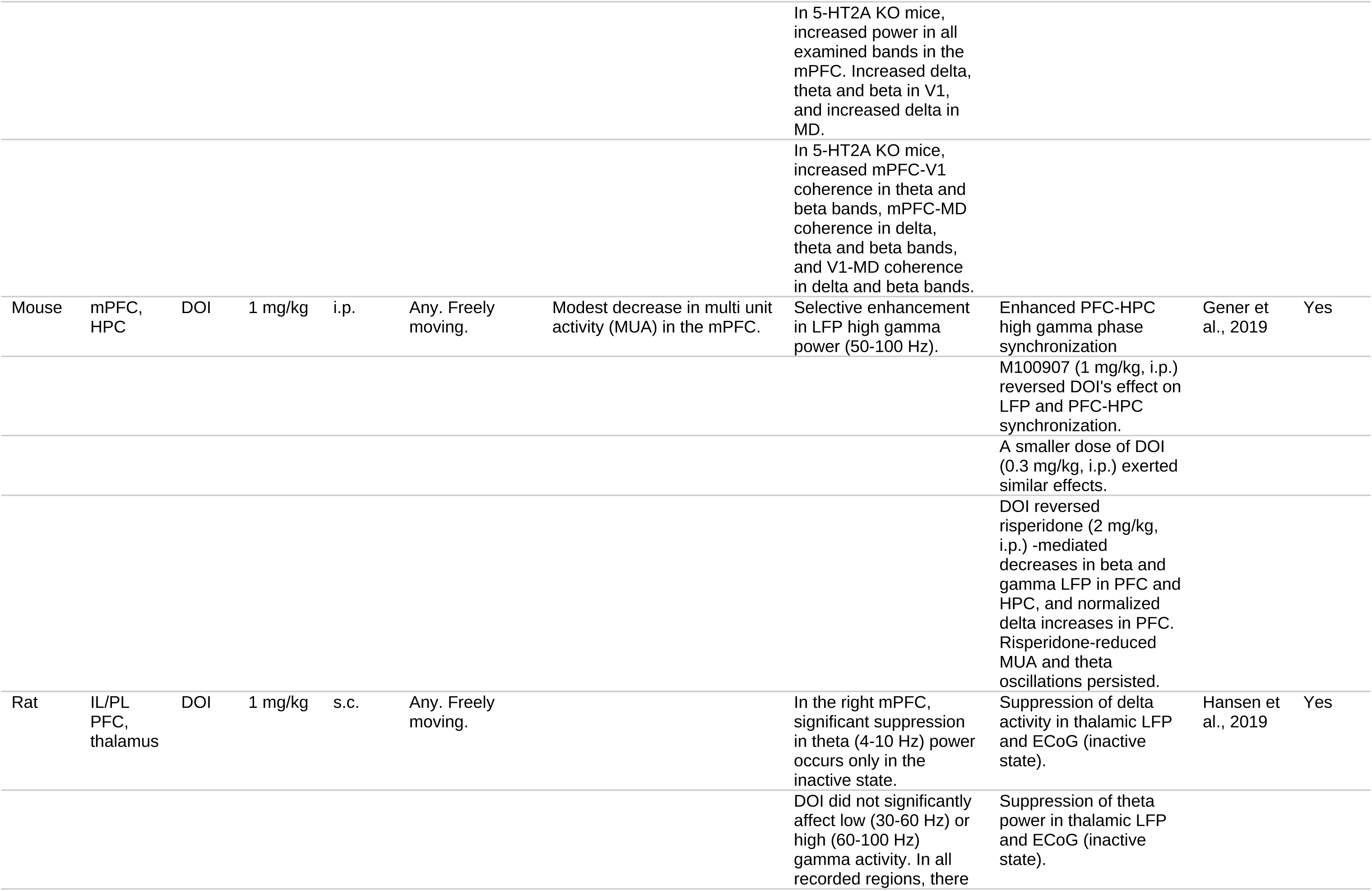
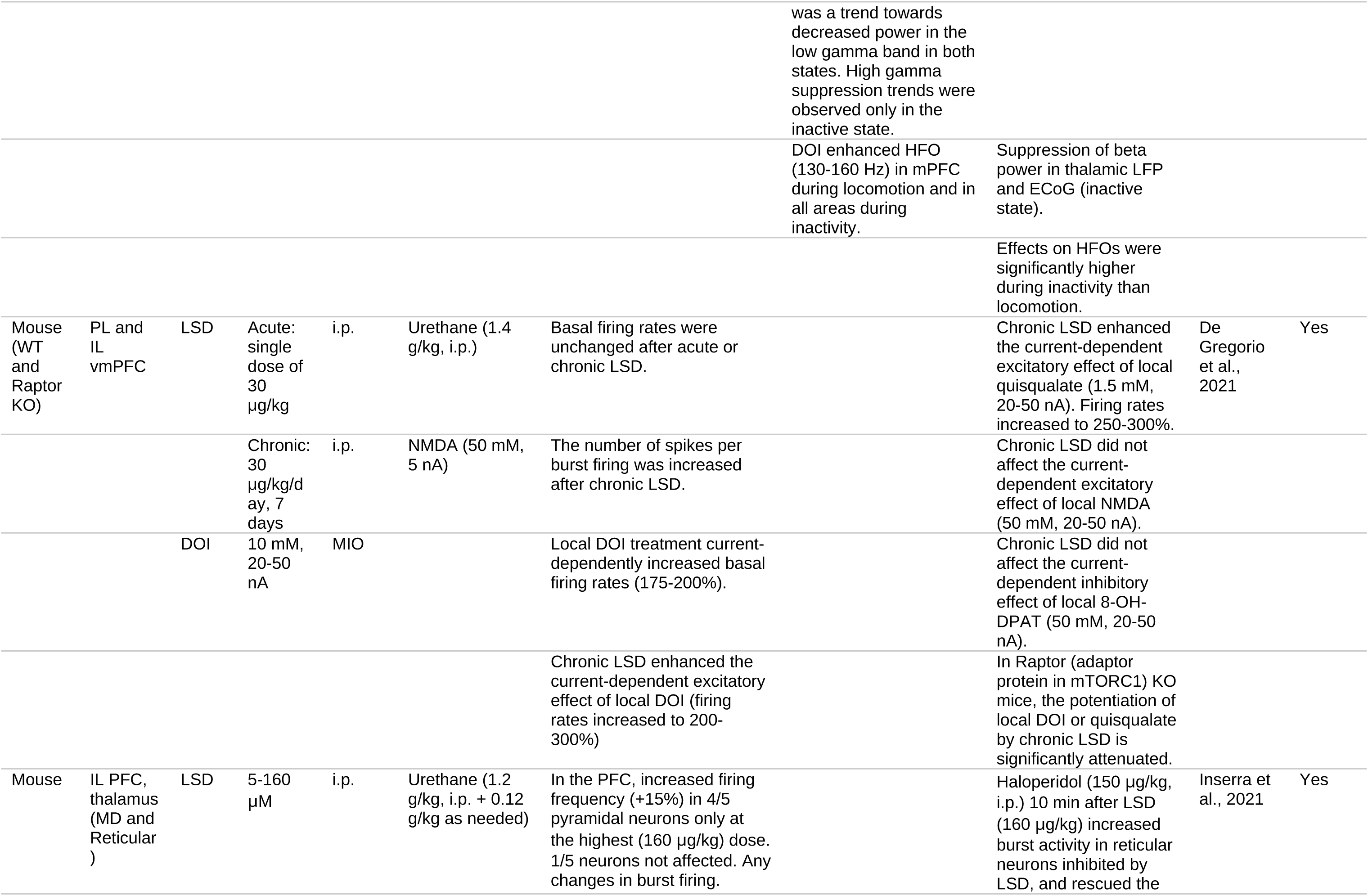
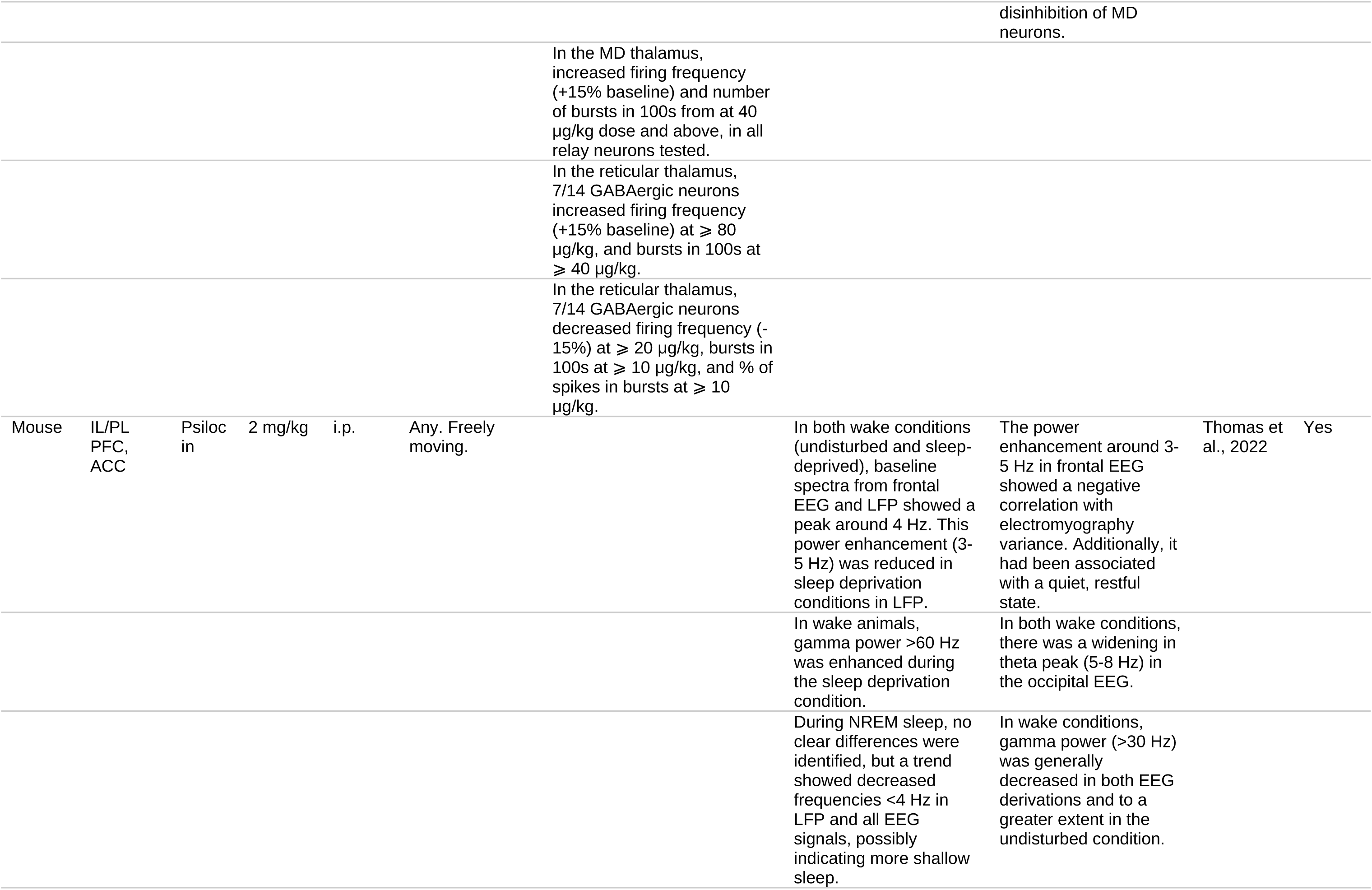
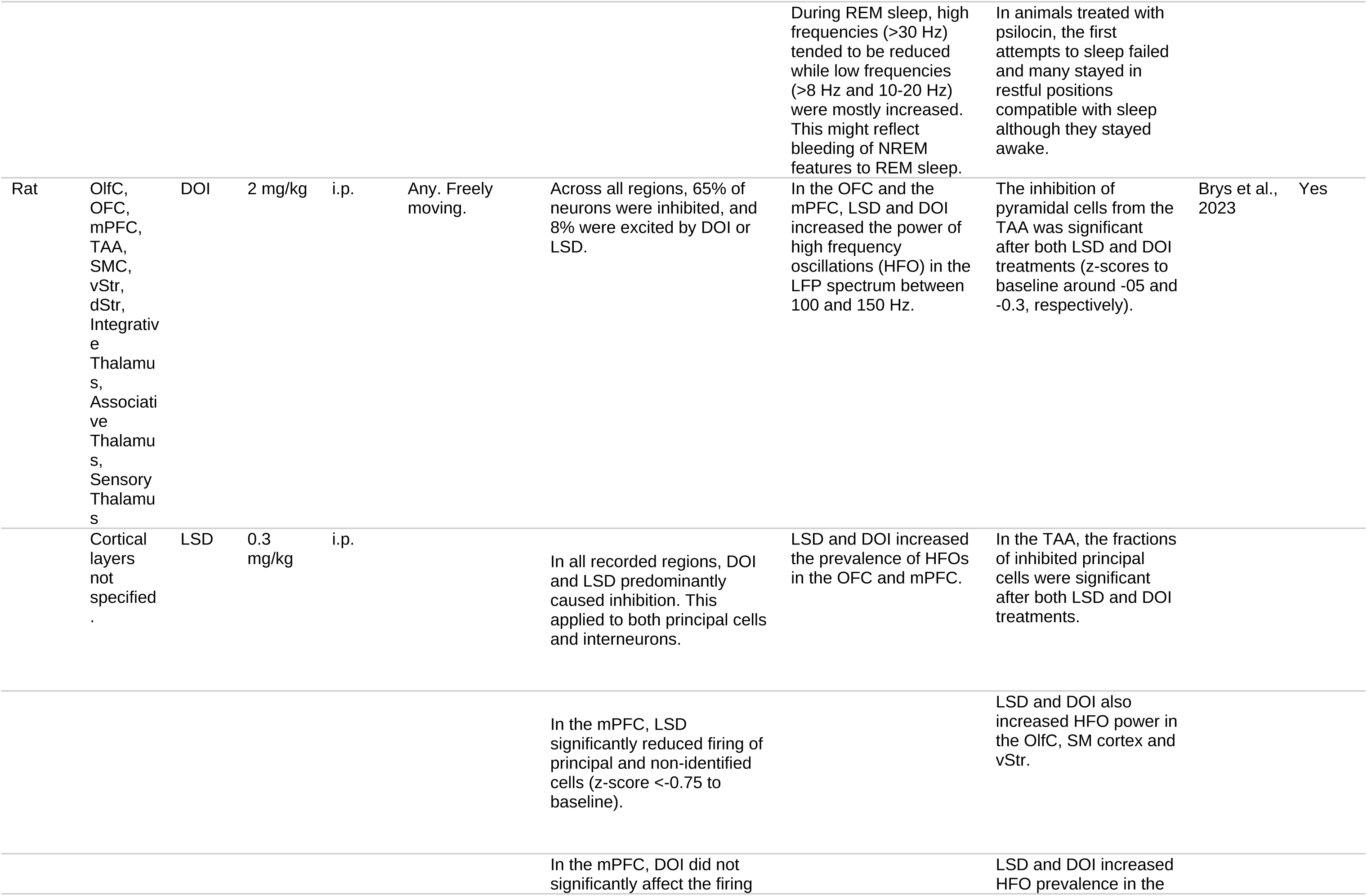
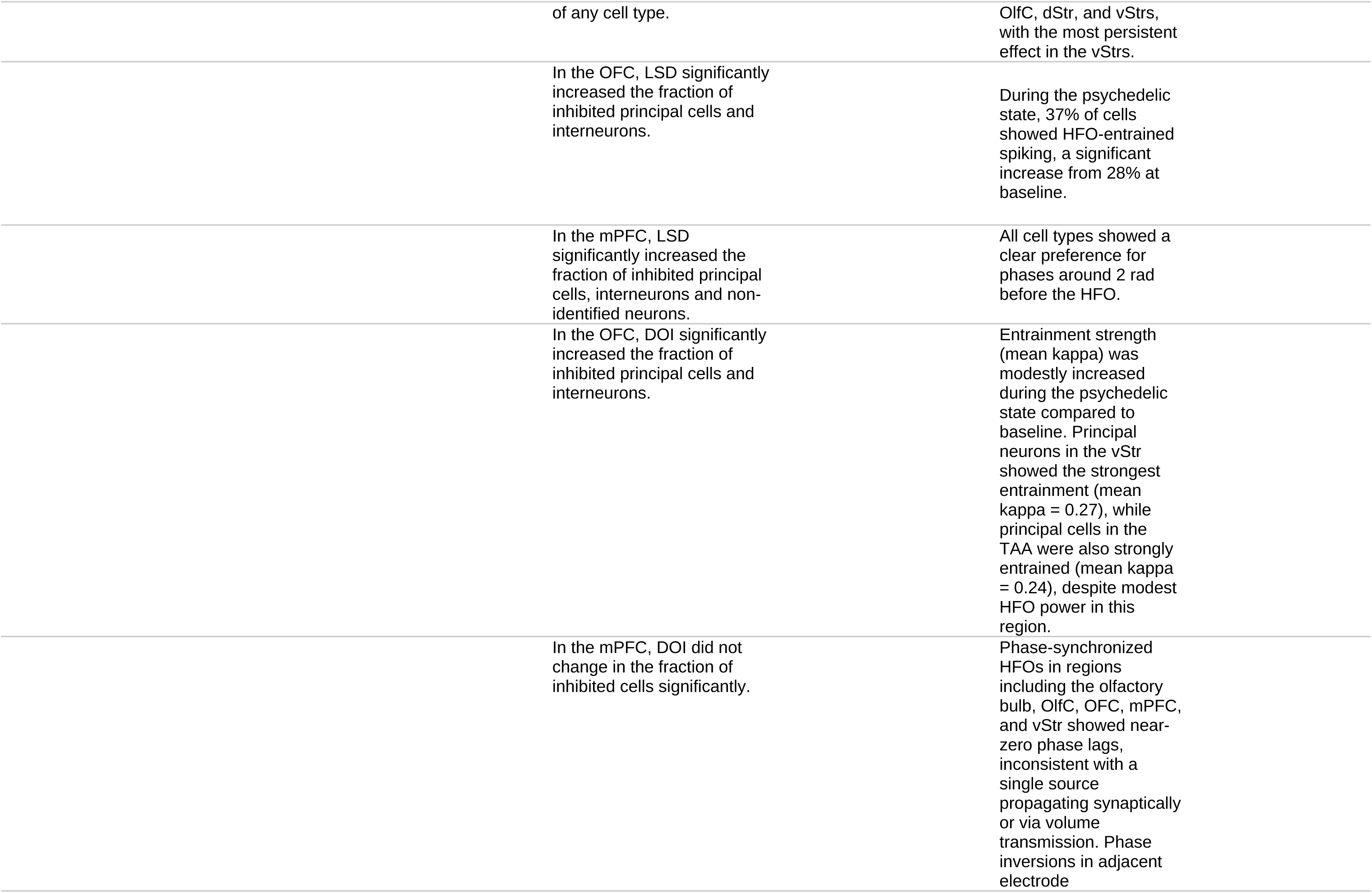
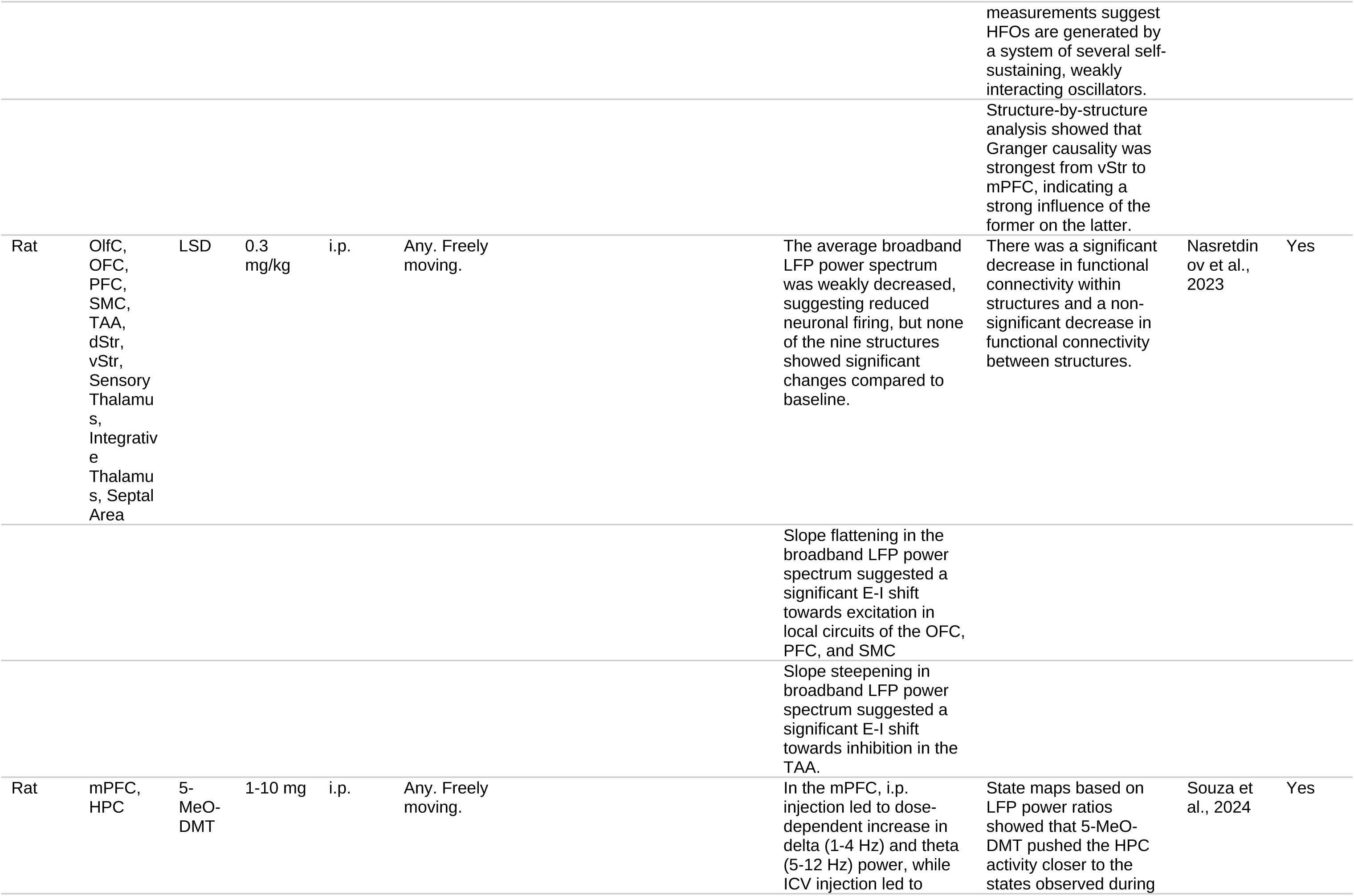
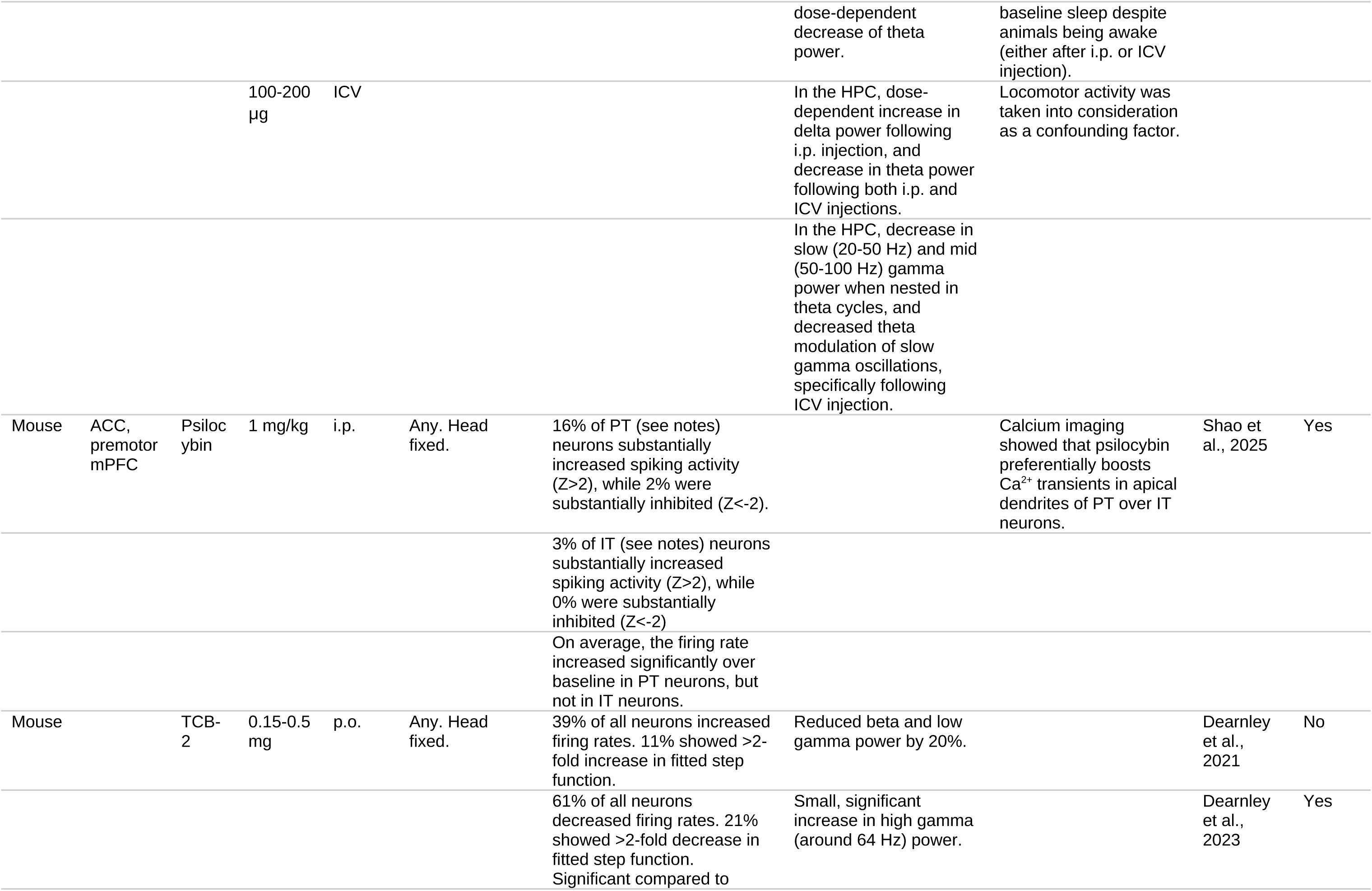
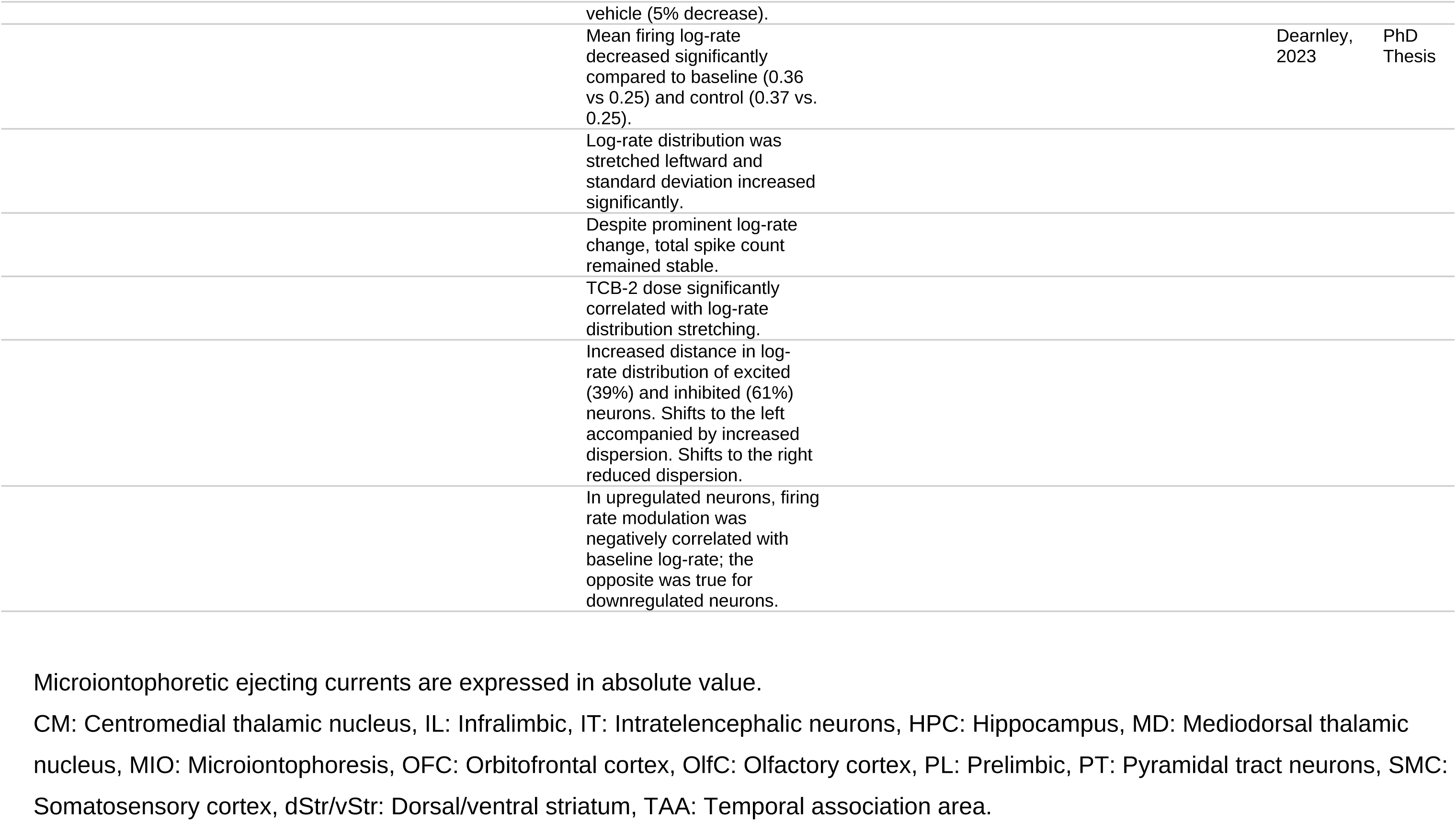
*In vivo* studies.

#### 3.2.1 Local Application

Microiontophoretic DOI delivery to the frontal cortex of rats, mice, and guinea pigs consistently induces current-dependent inhibition of both spontaneous and quisqualate (an AMPAR agonist) -inducased firing in L5/6p neurons from animals anesthetized with chloral hydrate (Ashby et al., 1989; Ashby & Wang, 1990; Bambico et al., 2010; Bergqvist et al., 1999; El Mansari & Blier, 1997, 2005; Rueter & Blier, 1999; Reuter et al., 2000). However, at lower ejecting currents (0.5 nA), DOI exerted a facilitatory effect on glutamate-induced firing in the mPFC, increasing firing rates in a fraction of the cells tested, whereas higher currents (>10 nA) suppressed all cells (Ashby et al., 1989; Ashby et al., 1994). Ritanserin and 5-HT_1A_ activation prevented the excitatory effect at low ejecting currents (Ashby et al., 1994). This biphasic modulation aligns with *in vitro* results from Arvanov et al. (1999a).

In the mPFC, DOI’s inhibitory action was unaffected by MgCl₂ (which prevents neurotransmitter release) but was blocked by several atypical neuroleptics, such as clozapine, ritanserin or risperidone, suggesting a direct, 5-HT_2A_-mediated mechanism (Ashby et al., 1989; Ashby and Wang, 1990; Bergqvist et al., 1999; Reuter and Blier, 1999). In contrast, in the OFC, DOI’s inhibition remained unaffected by atypical neuroleptics (Bergqvist et al., 1999; El Mansari and Blier, 1997), and by bicuculline (a GABA_A_ antagonist) (Bergqvist et al., 1999), but was blocked by the selective 5-HT_2A_ antagonist M100907 (Reuter et al., 2000). Intriguingly, knocking out 5-HT_2C_ receptor did not affect DOI’s inhibitory effect in the OFC, but abolished M100907’s blocking action (Reuter et al., 2000).

In mice, knocking out the fatty acid amide hydrolase (FAAH) significantly reduced DOI’s inhibitory effects, suggesting endocannabinoids play a modulatory role (Bambico et al., 2010).

However, one study presented conflicting results in mice, reporting a current-dependent increase in the firing rates of L5p following DOI application (20–50 nA) (De Gregorio et al., 2021). The discrepancy may stem from methodological differences: the use of urethane anesthesia, rather than chloral hydrate, and the use of NMDA to activate neurons within their physiological firing rates, rather than quisqualate. The excitatory response to local DOI administration was further enhanced after chronic systemic LSD treatment, an effect significantly attenuated by Raptor KO (De Gregorio et al., 2021). Raptor, an adaptor protein involved in mTORC1 signaling, is linked to neural plasticity, suggesting that chronic LSD effects may involve mTOR-dependent synaptogenesis (Ly et al., 2018).

Local DOI delivery into the mPFC by microdialysis significantly increased 5-HT release. This effect can be potentiated by DAMGO (a μ-opioid receptor agonist) and LY-379268 (a mGlu II receptor agonist) (Puig et al., 2003), raising the question of how endogenous 5-HT receptor activation contributes to DOI’s effects.

#### 3.2.2 Systemic Administration in Anesthetized Animals

##### 3.2.2.1 Firing activity

In chloral hydrate-anesthetized rats, systemic DOI administration produced a two-fold increase in the average firing rate of mPFC L5p neurons projecting either to the dorsal raphe or ventral tegmental area. About 40% of these neurons showed nearly quintupled firing rates, while 30% displayed a tenfold reduction from baseline activity. At the single neuron level, firing rate increases can reach up to 1525%, with the lower average rate attributed to the low baseline firing rates of some neurons. No significant differences in baseline firing rates could be observed between projecting L5p neurons that are upregulated, downregulated, or unaffected by DOI (Puig et al., 2003).

In neurons upregulated by DOI, burst firing increased concurrently, whereas in those where firing was downregulated, burst firing decreased (Celada et al., 2008). In most excited neurons, the firing rate increase was abolished by M100907 and LY-379268. In inhibited neurons, M100907 reversed the firing rate decrease in all cases. These findings suggest that the increase in firing rates is primarily mediated by 5-HT_2A_ agonism and glutamate release, yet 5-HT_2A_ activation also leads to neuronal inhibition in a subset of L5p neurons. Notably, M100907 remained effective in rats with thalamic lesions, indicating that firing rate modulation is mediated by non-thalamic 5-HT_2A_ receptors (Puig et al., 2003). The GABA_A_ antagonist picrotoxin also reversed systemic DOI’s inhibitory effects in the rat mPFC (Puig et al., 2003), contrasting with Bergqvist et al. (1999), who observed no reversal of local DOI’s inhibition by bicuculline in the rat OFC.

A combination of 5-MeO-DMT and the monoamine oxidase inhibitor (MAOI) clorgyline (loosely mimicking ayahuasca) produced similar effects: firing rates increased in half of the L5p neurons by an average of 406%, while 35% of neurons decreased their firing to 31% of baseline, resulting in an average net increase of 215% (Riga et al., 2014). Burst firing also increased. Both clozapine and haloperidol reversed the effects on firing rates and bursting (Riga et al., 2014).

While DOI does not activate 5-HT_1A_ receptor, that has inhibitory effects on L5p neurons (Nichols, 2016), 5-MeO-DMT stands out among psychedelics due to its higher affinity for 5-HT_1A_ than for 5-HT_2A_ (Puigseslloses et al., 2024). Therefore, it is surprising that DOI reduced L5p firing to a greater extent than 5-MeO-DMT (tenfold vs. two thirds decrease), and that almost the same proportion of cells was inhibited (30% vs. 35%) (Puig et al., 2003; Riga et al., 2014). These findings further support that 5-HT_2A_ is involved in the inhibitory action of psychedelics.

An acute dose of LSD had no significant effect on firing rates. Chronic LSD administration increased bursting activity in L5p neurons, yet it did not increase firing rates. Additionally, chronic LSD potentiated the excitatory response to quisqualate, but not to NMDA, nor the inhibitory action of the 5-HT_1A_ agonist 8-OH-DPAT. As noted earlier, the effects of chronic LSD are attenuated by Raptor KO and may be attributed to dendritic and synaptic plasticity (De Gregorio et al., 2021).

Despite the increase in firing rates elicited by 5-MeO-DMT, the BOLD signal in the mPFC and the primary visual cortex (V1) was significantly reduced. This discrepancy between electrophysiological and fMRI recordings could not be explained by alterations in blood pO₂ or pCO₂ (Riga et al., 2014). Additional studies have observed decreased BOLD signals in rats and humans following psychedelic administration (Carhart-Harris et al., 2012; Ghaw et al., 2024). Although firing rates are increased in L5p projecting neurons (Puig et al., 2003; Riga et al., 2014; Shao et al., 2025), psychedelics exert an overall inhibitory effect on neuronal firing (Brys et al., 2023; Dearnley et al., 2021). Another possible explanation for this discrepancy is that psychedelics may reduce aerobic metabolism, as suggested by the downregulation of mitochondrial genes and induction of resistance to hypoxia and ischemia in tissues (Nardai et al., 2020; Szabo et al., 2016; Szabó et al., 2021; Zhou et al., 2025).

Under ketamine anesthesia, LSD (30-60 μg/kg, i.p.) did not change PFC firing rates (Godbout et al., 1991). Moreover, it markedly prolonged the inhibition of PFC neurons following electrical stimulation of the ventral tegmental area, which could be prevented using a 5-HT_2A_ antagonist (Godbout et al., 1991). Congruent with this finding, under urethane anesthesia, LSD could increased mPFC firing frequency only at the highest dose tested (160 μg/kg, i.p.) (Inserra et al., 2021). No significant change in burst firing was observed. By contrast, at lower doses (10–80 μg/kg, i.p.), all tested neurons in the mediodorsal nucleus (MD) of the thalamus increased both their firing frequency and bursting, while in the reticular nucleus, half of the neurons were excited and the other half inhibited. Haloperidol was able to rescue the inhibition in the reticular nucleus (Inserra et al., 2021).

##### 3.2.2.2 Local Field Potentials

Both DOI and 5-MeO-DMT reduced low-frequency oscillations (LFO, <4 Hz) within mPFC local field potentials (LFP), with 5-MeO-DMT additionally suppressing LFO in V1 and other unimodal cortices (Celada et al., 2008; Riga et al., 2014, 2016). This suppression occurred independently of whether the firing rate of a recorded neuron was increased or decreased (Riga et al., 2014). Notably, the proportion of spikes occurring during the active phase of LFO, defined as the portion below its mean voltage value, decreased indicating a decoupling of oscillatory activity from spike timing (Celada et al., 2008). The LFO reduction was partially or completely blocked by 5-HT_2A_ antagonists, atypical neuroleptics and 5-HT_2A_ KO (Celada et al., 2008; Riga et al., 2014, 2016). For 5-MeO-DMT specifically, the 5-HT_1A_ antagonist WAY-100635 and the mGlu II agonist LY-379268 also blocked LFO suppression (Riga et al., 2014, 2016).

#### 3.2.3 Systemic Administration in Awake Animals

##### 3.2.3.1 Firing activity

Awake animal studies mostly report inhibitory or neutral effects of psychedelics on neuronal firing (Brys et al., 2023; Dearnley et al., 2021; Gener et al., 2019; Wood et al., 2012).

The most comprehensive electrophysiological study to date has recorded 3,612 mPFC neurons in mice following TCB-2 administration, a highly 5-HT_2A_-selective psychedelic (Dearnley et al., 2021). [For a peer-reviewed version of this work, but focusing on the comparison of TCB-2’s effects to arousal, see (Dearnley et al., 2023). For the complete PhD thesis, see (Dearnley, 2023)]. However, this study did not isolate L5p neurons from other neural types. Of all neurons, 11% were significantly excited, and 21% were inhibited. Although the mean logarithmic firing rate decreased significantly, the average firing rate and the total number of spikes remained stable, due to a redistribution in spiking activity. The distribution of decreased logarithmic rates shifted leftward and became more dispersed, while that of increased log-rates became more condensed to the right (Dearnley et al., 2021). This pattern, where inhibition of many neurons offsets excitation of fewer fast-spiking (typically non-pyramidal) cells, challenges the simplistic view of psychedelics as uniformly excitatory in the mPFC, which is at the basis of some of the leading hypotheses about psychedelic action.

These compensatory dynamics may be masked by firing rate normalization. Normalized firing rates of excitatory and inhibitory cells were significantly decreased by DOI and LSD in many cortical and subcortical regions of freely moving rats, including the mPFC L5p neurons, with the most significant reductions occurring in the temporal association area (Brys et al., 2023; Wood et al., 2012).

However, 16% of mouse mPFC L5p neurons projecting to the pons substantially increased their spiking activity after psilocybin injection, while only 2% are substantially inhibited, with the average firing rate being increased (Shao et al., 2025). In contrast, only 3% of L5p neurons projecting to the caudate putamen (i.e., intratelencephalic neurons) were excited, and none were inhibited. Congruently, *in vivo* calcium imaging revealed significant increases in calcium transients in projecting, but not intratelencephalic neurons (Shao et al., 2025).

The finding that psychedelics increase the average firing rate of L5p projecting neurons is in line with the findings of Puig et al. (2003) and Riga et al. (2014) in anesthetized animals. Nevertheless, Shao et al. (2025) may have underestimated the proportion of inhibited neurons. They apply a criterion of |Z| > 2 to classify substantial increases or decreases in firing rates. As described by Dearnley et al. (2021), the distribution of excited neurons is concentrated toward the right, while the distribution of inhibited neurons is spread toward the left. Consequently, Shao et al. (2025) should have observed a much higher number of excited neurons in the right tail (Z>2) than in the left tail (Z<-2) of the distribution curve.

As for population-level firing, a modest decrease in multi-unit activity was seen in the mouse prelimbic frontal cortex following DOI administration (Gener et al., 2019).

##### 3.2.3.2 Local Field Potentials

LSD and DOI increased the power and prevalence of high-frequency oscillations (HFOs) in the 100–150 Hz range within LFP recorded from the rat mPFC and other brain regions (Brys et al., 2023). The increase was significantly greater when the animals are at rest (Hansen et al., 2019). The synchronized HFOs activity across the mPFC, OFC, and ventral striatum exhibited near-zero phase lags. Granger causality analyses indicated the ventral striatum exerts the strongest influence on the mPFC (Brys et al., 2023). At lower frequency bands, 5-MeO-DMT increases mPFC-V1 coherence in the beta band (10-30 Hz), mPFC-MD coherence in the theta (4-10 Hz) and beta bands, and V1-MD coherence in the beta band (Riga et al., 2018).

The effects on gamma frequency oscillations show more variability across studies. DOI administration decreases both low (30–55 Hz) and high (55–80 Hz) gamma power in rat OFC, with the high gamma reduction diminishing at higher doses (Wood et al., 2012). In the ACC, DOI reduces both gamma sub-bands at increasing doses (Wood et al., 2012). Other studies report more modest effects in rats, with non-significant decreases in mPFC low gamma power that extend to the high gamma band when animals are at rest (Hansen et al., 2019). In turn, psilocin increases high gamma power (>60 Hz) in the mPFC and ACC in sleep-deprived mice (Thomas et al., 2022), while DOI increases mPFC high gamma power (50–100 Hz) in undisturbed mice, enhancing synchronicity between the mPFC and hippocampus, effects that are reversed by antagonizing 5-HT_2A_ (Gener et al., 2019). As for 5-MeO-DMT, it increases gamma power (30–80 Hz) in the mPFC via both 5-HT_2A_ and 5-HT_1A_ activation (Riga et al., 2018).

In lower-frequency bands, DOI decreases theta power (4–10 Hz) in the mPFC of resting mice (Hansen et al., 2019). In contrast, a psilocin injection given at the typical sleep onset time enhanced peak at 4 Hz only when animals were allowed to rest (Thomas et al., 2022). As for 5-MeO-DMT, i.p. injection led to a dose-dependent increase in delta (1–4 Hz) and theta (5–12 Hz) power, while intracerebroventricular (ICV) injection led to a dose-dependent decrease in theta power (Souza et al., 2024).

The correlations between oscillatory activity and spike output also show inconsistent alterations. DOI reduces the correlation between gamma oscillations and spike timing in the rat OFC and ACC (Wood et al., 2012). This result aligns with the findings of Celada et al. (2008) in mPFC LFO of anesthetized rats. However, LSD and DOI exerted a modest increase in the entrainment strength of spikes by HFOs in the rat mPFC, and increased the proportion of neurons exhibiting entrainment. The ventral striatum, which appears to drive HFO synchronicity between regions, shows particularly strong HFO-spike entrainment (Brys et al., 2023).

At the network level, LSD reduces broadband LFP power across multiple regions, including OFC and mPFC, suggesting a shift toward reduced firing (Nasretdinov et al., 2023), which is consistent with findings from Brys et al. (2023) and Dearnley et al. (2021). Furthermore, changes in LFP spectral slopes suggested an E/I balance shift toward excitation in local PFC circuits, thus, a decoupling between the network’s E/I state and spike output (Nasretdinov et al., 2023). In the temporal association area, however, there is a congruent E/I shift towards inhibition, decrease in firing activity, high HFO entrainment strength and low HFO prevalence (Brys et al., 2023; Nasretdinov et al., 2023).

As a final note, some studies have reported the emergence of sleep-like electrophysiological features in awake animals (Souza et al., 2024; Thomas et al., 2022).

## 4. Discussion

The aim of this review work was to facilitate a comprehensive view of electrophysiological data about the action of psychedelic drugs. This was needed in order to complement data from neuroimaging studies with more precise physiological experimentation. Our main interest was to challenge one prevailing assumption in psychedelic science: that psychedelics increase neuronal excitability in L5p neurons. We focused on the frontal cortex since it is the brain region which has been most studied, and reviewed data from *in vitro* experimentation, which afford a precise control of the recordings, as well as from *in vivo* assays that reproduce the complexity of a living brain.

### 4.1 Summary

Psychedelics exert a complex modulatory influence on L5p neurons, combining excitatory and inhibitory effects.

*In vitro*, they induce a mild but consistent depolarization of the RMP (Araneda & Andrade, 1991; Arvanov et al., 1999a; Ekins et al., 2023; Lambe & Aghajanian, 2006; Spindle & Thomas, 2014; Tanaka & North, 1993) while suppressing NMDA-evoked EPSCs/EPSPs (Arvanov et al., 1999a, 1999b). They also trigger AMPAR internalization, further dampening excitatory transmission (Berthoux et al., 2019), though a subset of L5p neurons exhibit potentiated AMPA-induced currents (Arvanov et al., 1999a). Despite these suppressive effects, psychedelics shift the E/I balance toward excitation, producing late, prolonged eEPSCs/eEPSPs that correspond to up states (Aghajanian & Marek, 1999; Marek & Ramos, 2018; Lambe & Aghajanian, 2006, 2007; Stutzmann et al., 2001).

Firing responses vary: some studies report marked increases in evoked and spontaneous L5p activity (Araneda & Andrade, 1991; Schmitz et al., 2022), while others observe concentration-dependent suppression (Ekins et al., 2023; Wang et al., 2025) or non-significant changes (Lambe & Aghajanian, 2006). Similarly, spike frequency adaptation may decrease (Araneda & Andrade, 1991; Arvanov et al., 1999a) or increase (Ekins et al., 2023). Notably, psychedelics consistently reduce AHP (Araneda & Andrade, 1991; Arvanov et al., 1999a; Villalobos et al., 2005).

*In vivo*, microiontophoretic application of DOI inhibits L5p firing (Ashby et al., 1989; Ashby & Wang, 1990; Bambico et al., 2010; Bergqvist et al., 1999; El Mansari & Blier, 1997, 2005; Reuter & Blier, 1999), whereas systemic administration yields mixed excitatory, inhibitory, or neutral effects across populations (Celada et al., 2008; Godbout et al., 1991; Inserra et al., 2021; Puig et al., 2003; Riga et al., 2014; Shao et al., 2025). The dominant outcome is net inhibition of L5p cells (Wood et al., 2012), as well as of excitatory and inhibitory neurons more generally (Brys et al., 2023). However, this suppression is offset by increased firing in a subset of fast-spiking neurons, preserving overall mPFC spike output (Dearnley et al., 2021). See also Béïque et al. (2007); Martin & Nichols (2016)

LFP effects are inconsistent. However, most studies report reduced power in low-gamma and lower frequencies (Celada et al., 2008; Hansen et al., 2019; Riga et al., 2014, 2016; Souza et al., 2024) but elevated high-gamma and higher-frequency activity (Brys et al., 2023; Gener et al., 2019; Riga et al., 2018; Thomas et al., 2022). Some LFPs also show enhanced cross-regional synchrony (Brys et al., 2023; Gener et al., 2019; Riga et al., 2018). While LFP-spike coupling typically weakens in low-frequency and gamma bands (Celada et al., 2008; Wood et al., 2012), some report stronger HFO-spike entrainment (Brys et al., 2023).

24 hours post-administration, studies note increased mEPSCs, eEPSCs, and burst firing (De Gregorio et al., 2021; Ly et al., 2018; Shao et al., 2021), likely reflecting synaptic and dendritic plasticity rather than direct electrophysiological modulation.

### 4.2 Differential Dendritic Modulation by Psychedelics

Anatomically, L1 in the PFC receives long-range fibers from higher-order thalamic nuclei projecting to the apical dendritic tufts of L5p neurons (Anastasiades et al., 2021), as well as cortico-cortical projections (Mitchell & Cauller, 2001). L2/3 is traversed by the apical dendritic trunks of L5p cells, and the most superficial part of L5 (L5a) contains oblique dendrites. The soma and proximal dendrites of L5p neurons are located deeper in L5 (Larkum, 2013; Squire et al., 2012). Functionally, L5p neurons can be divided into three compartments: (1) a basal compartment (soma-proximal dendrites), (2) an apical compartment (tuft dendrites), and (3) a coupling compartment (apical trunk and oblique dendrites) (Aru et al., 2020; Suzuki & Larkum, 2020).

According to Dendritic Integration Theory (Aru et al., 2020; Bachmann et al., 2020), the basal compartment receives feedforward sensory input, whereas the apical compartment integrates higher-order thalamo-cortical and cortico-cortical contextual signals. These streams converge in the coupling compartment, which is modulated by higher-order thalamic nuclei (Aru et al., 2020; Bachmann et al., 2020). During wakefulness, apical and basal compartments remain coupled, enabling sensory-context integration. Under general anesthesia, this coupling is disrupted, isolating the two streams (Aru et al., 2020; Suzuki & Larkum, 2020).

Psychedelics appear to modulate distinct compartments of L5p neurons in a layer-specific manner. Studies stimulating L1 report increased eEPSPs/eEPSCs (Aghajanian & Marek, 1997; Barre et al., 2016), whereas those stimulating L5 or the forceps minor observe either unchanged eEPSPs or reduced eEPSCs/eEPSPs (Arvanov et al., 1999a, 1999b; Tanaka & North, 1993). Therefore, psychedelics enhance electrically- and NMDA-evoked EPSCs/EPSPs in the apical compartment (Aghajanian & Marek, 1997; Barre et al., 2016), while suppressing NMDAR-mediated eEPSCs/eEPSPs more broadly (Arvanov et al., 1999a). This suppressive effect is particularly pronounced with 5-HT1A agonists like LSD or 5-HT itself (Arvanov et al., 1999b). Given that 5-HT1A is an inhibitory, Gαi/o-coupled receptor which expression is enriched in the basal compartment and axon hillock (Czyrak et al., 2003), these findings suggest that psychedelics, especially those with significant 5-HT1A activity like 5-MeO-DMT (Puigseslloses et al., 2024), bias L5p neuron excitability toward the apical compartment.

Regarding the dendritic coupling region, it is noteworthy that this area concentrates the highest expression of 5-HT_2A_ proteins in the neocortex (Chiu et al., 2023; Weber & Andrade, 2010). Backpropagating Na^+^ action potentials facilitate the firing of dendritic Ca^2+^ spikes in the L5p dendritic trunks, which in turn promotes axonal burst firing (Larkum et al., 1999). Psychedelics are known to increase burst firing in L5p neurons (Celada et al., 2008; Riga et al., 2014), and ROB is generated intrinsically via a direct, 5-HT_2A_-mediated mechanism (Spindle & Thomas, 2014). Furthermore, psychedelics can promote up states during slow (<1 Hz), rhythmic stimulation in the cortical midlayers (Lambe & Aghajanian, 2006, 2007). Altogether, psychedelics seem to enhance the excitability of the coupling region, thereby increasing tuft-to-soma signal transmission. This effect directly opposes that of general anesthetics like isoflurane, urethane and ketamine/xylazine (Suzuki & Larkum, 2020); indeed, recent work shows that DOI reverses the effects of propofol and isoflurane (Huels et al., 2025). Intriguingly, ketamine is noted for its psychedelic-like effects. Studies such as those of Brys et al. (2023) highlight how its mechanisms might converge with those of serotonergic psychedelics (see also Ali et al., 2020), despite their seemingly opposite actions on L5p coupling. The characterization of convergent and divergent mechanisms of serotonergic psychedelics and psychedelic-like anesthetics remains a topic for future research.

As mentioned above, psychedelics facilitate eEPSCs following L1 stimulation, but not stimulation of the forceps minor (Aghajanian & Marek, 1997; Arvanov et al., 1999a, 199b; Barre et al., 2016; Tanaka and North, 1993). While the forceps minor consists of interhemispheric cortico-cortical (callosal) fibers, L1 receives a dense convergence of cortico-cortical fibers and projections from higher-order thalamic nuclei, such as the MD nucleus (Anastasiades et al., 2021). The differential effect of L1 vs. forceps minor stimulation suggests a selective boosting of thalamo-cortical and/or ipsilateral feedback, rather than interhemispheric callosal inputs. Congruently, psychedelics reduce the amplitudes of EPSCs evoked by callosal stimulation (Troca-Marín & Geijo-Barrientos, 2010). Furthermore, psychedelics increase firing frequency and bursting in the MD nucleus, while enhancing mPFC-MD coherence in the theta (4–10 Hz) band (Inserra et al., 2021; Riga et al., 2018).

Apical hyperexcitability and increase tuft-to-soma signal propagation, combined with basal hypoexcitability, amplifies apical influence over L5p output, termed apical drive (Phillips et al., 2025). Thus, by increasing apical drive the psychedelic state may reflect contextual, top-down thalamo-cortical dominance over L5p activity. Some hypotheses attribute hallucinations to increased top-down processing (Corlett et al., 2019), aligning with psychedelics reducing sensory-evoked visual cortex activity (Heiss et al., 1973; Michaiel et al., 2019). Notably, these electrophysiological findings challenge hypotheses proposing either weakened top-down constraints (Carhart-Harris & Friston, 2019) or bottom-up sensory dominance (Vollenweider & Preller, 2020).

Several distinctions must be further clarified regarding the key differences between our proposal and prevailing models of psychedelic action, namely the CSTC hypothesis (Vollenweider & Preller, 2020). The CSTC model posits a primarily suppressive role for the thalamus, often described through the ‘sensory filter’ metaphor. According to this view, psychedelics inhibit GABAergic neurons in the thalamus, particularly within the thalamic reticular nucleus, thereby ‘unfiltering’ the arrival of bottom-up sensory stimuli to the cortex. This cortical hyperactivity then loops back to the thalamus via the striatum (Vollenweider & Preller, 2020).

In contrast, we argue that the thalamic nuclei central to psychedelic action are not inhibitory structures like the reticular nucleus, but higher-order excitatory nuclei, such as the MD nucleus. Supporting this view, Inserra et al. (2021) observed that while the effects of LSD on the reticular nucleus neurons were inconsistent (with roughly half being excited and half inhibited), LSD consistently increased the firing rate of every MD neuron recorded.

Furthermore, while the CSTC model emphasizes the relevance of the indirect cortico-striato-thalamic pathway, driven by IT neurons in Layer 5, we argue that the direct cortico-thalamic pathways driven by ET neurons are selectively excited by psychedelics. This mechanism has been empirically supported by Shao et al. (2025) and is further discussed in the section *Excitatory vs. Inhibitory Action of Psychedelics* below. This distinction is fundamental: rather than a failure of sensory filtering, we propose a gain-of-function in higher-order integration.

### 4.3 Molecular Mechanisms of Synaptic Modulation

Most *in vitro* studies have employed “low” concentrations (<3 μM) of DOB or DOI. However, the peak forebrain concentration of DOI following a systemic dose of 1 mg/kg, a standard dose used in in vivo electrophysiological studies (Table S2), reaches approximately 500 ng/g wet tissue weight, equivalent to ∼1.5 nM (De La Fuente Revenga et al., 2019). Phenylethylamines are particularly useful for isolating the core effects of psychedelics due to their higher selectivity for 5-HT_2A_ receptors compared to tryptamines and ergolines (Nichols, 2016). At these low concentrations, 5-HT_2A_-mediated effects follow a consistent temporal profile. Initially, DOB or DOI enhance eEPSPs/eEPSCs amplitudes by promoting presynaptic glutamate release (Arvanov et al., 1999a; Barre et al., 2016).

However, this potentiation undergoes significant desensitization within 15–20 minutes (Arvanov et al., 1999a). During this window, postsynaptic processes come into play. PKC drives AMPAR internalization (Berthoux et al., 2019), while CaM-KII suppresses NMDA-evoked currents (Arvanov et al., 1999a). Notably, in apical dendrites, PKC activity counters this inhibition by potentiating GluN2B-NMDAR (Barre et al., 2016), which may explain the sustained elevation of eEPSPs beyond the 20-minute mark (Aghajanian and Marek, 1997). In parallel, extrasynaptic GluN2B-NMDAR-dependent up states begin to emerge (Aghajanian & Marek, 1999; Lambe & Aghajanian, 2006, 2007; Stutzmann et al., 2001). Once established, these effects can persist for hours after washout (Arvanov et al., 1999a; Berthoux et al., 2019; Lambe & Aghajanian, 2006). To provide a schematic, integrated view of these dynamics, we developed Figures 2, 3 and 4.

**Figure 2:**
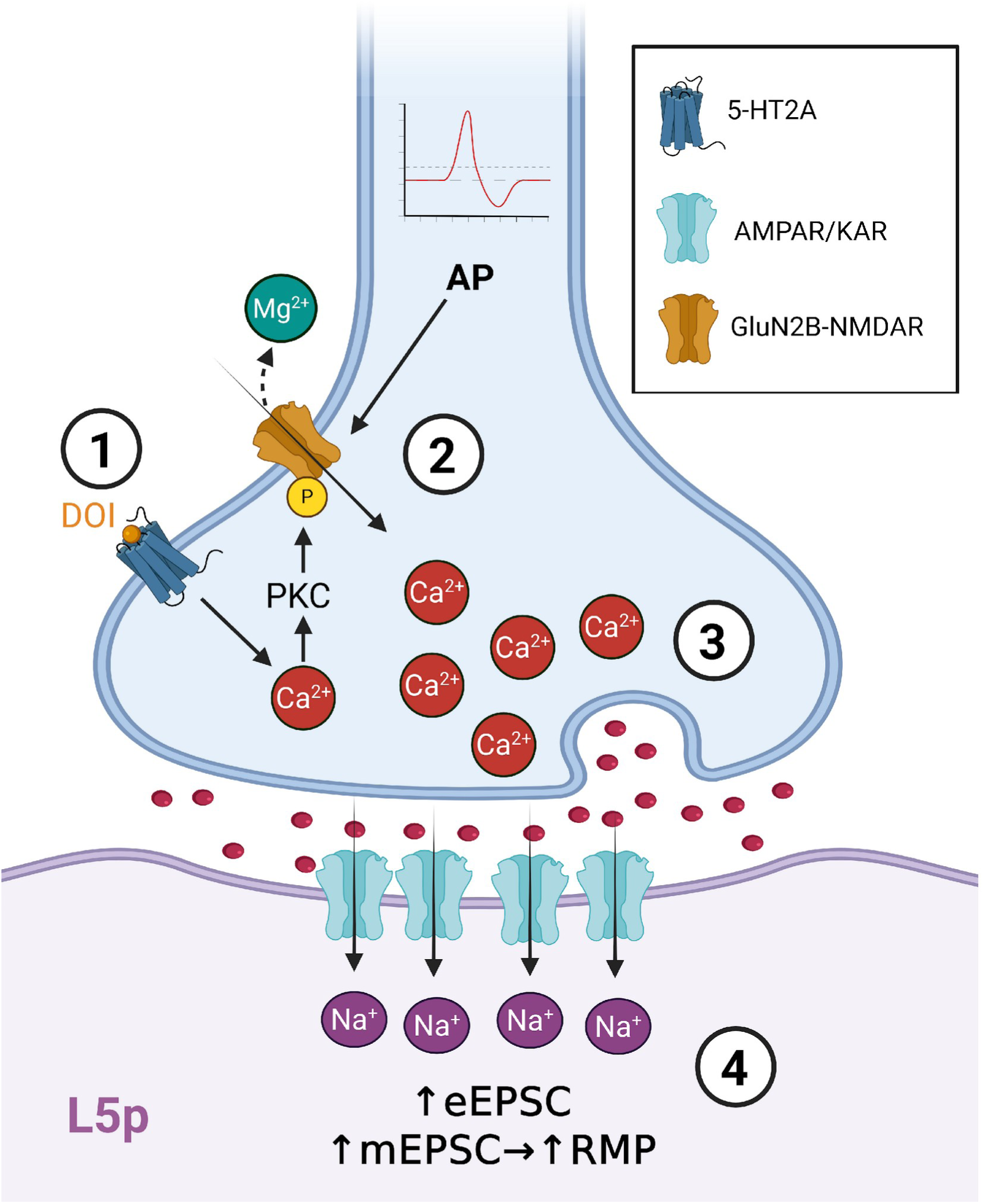
Early Response to DOI. Psychedelics lacking affinity for the 5-HT1A receptor can elicit an early excitatory response at low doses. Here, 2,5-dimethoxy-4-iodoamphetamine (DOI) is illustrated as a prototypical 5-HT2A-selective psychedelic. **1.** DOI binds to presynaptic 5-HT2A receptors, activating protein kinase C (PKC), which in turn phosphorylates GluN2B-containing NMDA receptors (GluN2B-NMDARs), enhancing their conductance. **2.** Upon the arrival of an action potential, the Mg²⁺ block is relieved from the NMDARs, allowing Ca²⁺ ions to enter the presynaptic terminal. **3.** Elevated presynaptic Ca²⁺ levels increase glutamate release (Red dots) into the synaptic cleft, activating postsynaptic AMPA and kainate receptors (AMPARs/KARs). Synchronous activation increases the amplitude of evoked excitatory postsynaptic currents (eEPSC), while asynchronous activation increases the frequency of miniature excitatory postsynaptic currents (mEPSC), contributing to a rise in the resting membrane potential (RMP).

**Figure 3:**
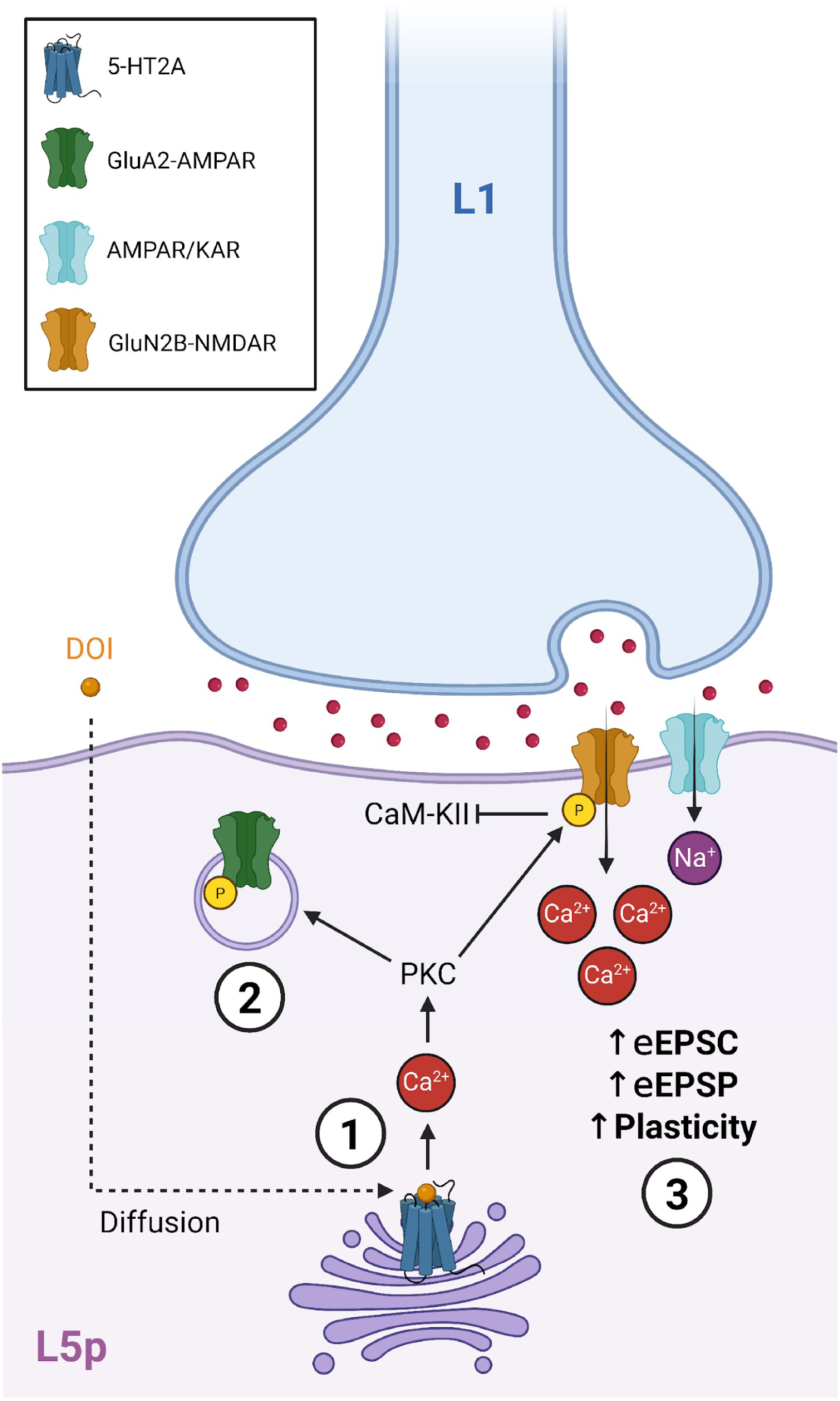
Late Response to DOI in Apical Tuft Dendrites. Psychedelics lacking affinity for the 5-HT1A receptor elicit a late excitatory response at low doses in the tuft dendrites of layer 5 pyramidal neurons. Here, 2,5-dimethoxy-4-iodoamphetamine (DOI) is illustrated as a prototypical 5-HT2A-selective psychedelic. **1.** DOI diffuses through the postsynaptic membrane, reaching intracellular 5-HT2A receptors, where activation recruits protein kinase C (PKC) via Ca²⁺-dependent signaling. **2.** PKC phosphorylates GluA2-containing AMPA receptors (GluA2-AMPARs), promoting their internalization and reducing AMPA-mediated transmission. **3.** At the same time, PKC phosphorylates GluN2B-containing NMDA receptors (GluN2B-NMDARs), enhancing their conductance and preventing modulation by Ca²⁺/calmodulin-dependent protein kinase II (CaM-KII). The activation of AMPA and kainate receptors (AMPARs/KAR) allows Ca²⁺ entry through GluN2B-NMDAR, boosting evoked excitatory postsynaptic currents (eEPSC), evoked excitatory postsynaptic potentials (eEPSPs), and promoting structural plasticity.

**Figure 4:**
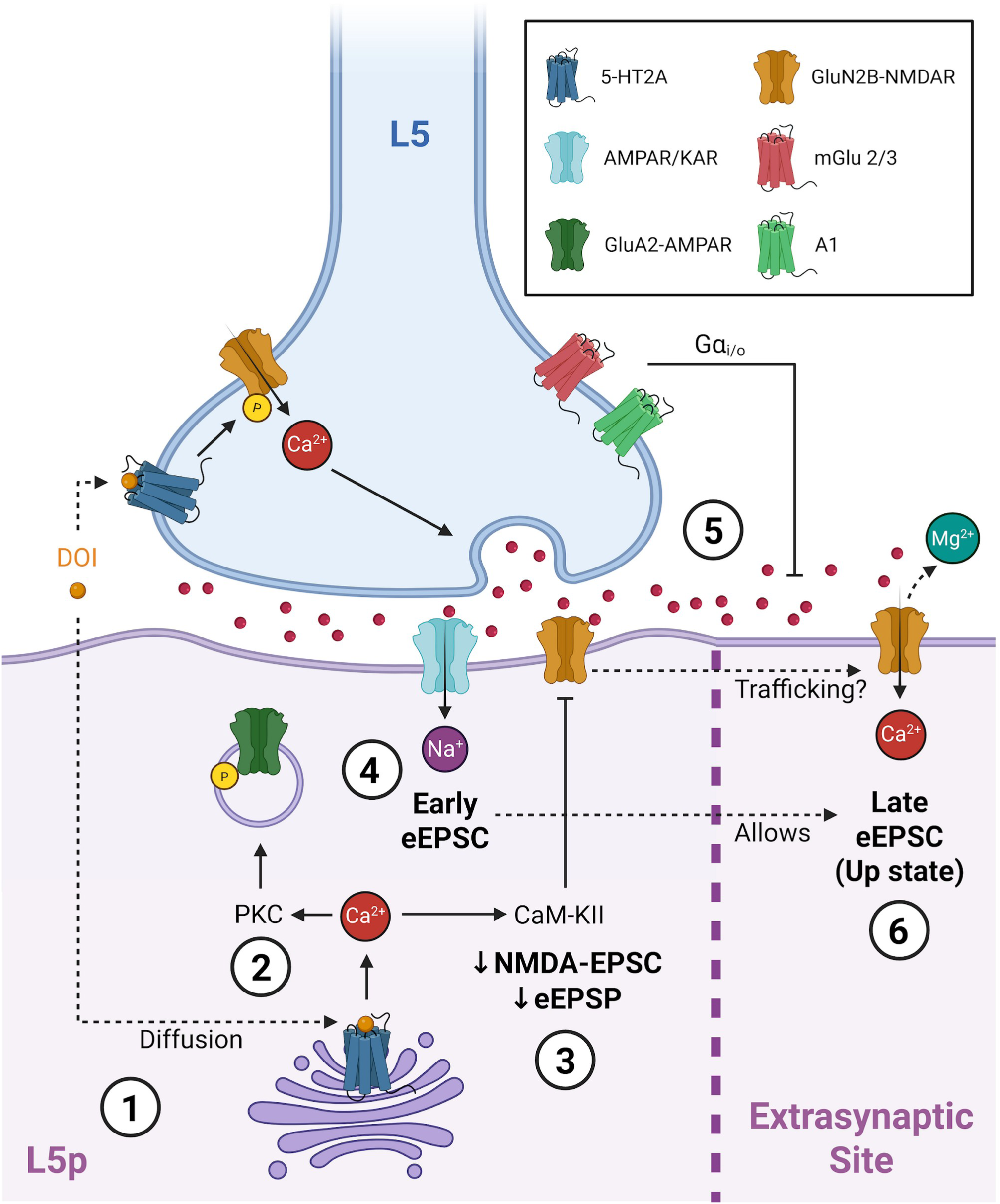
Late Response to DOI in Non-Apical Tuft Dendrites. Psychedelics lacking affinity for the 5-HT_1A_ receptor elicit a complex late response at low doses in basal dendrites of layer 5 pyramidal neurons. Here, 2,5-dimethoxy-4-iodoamphetamine (DOI) is illustrated as a prototypical 5-HT_2A_-selective psychedelic. 1. DOI diffuses through the postsynaptic membrane, reaching intracellular 5-HT2A receptors and triggering Ca²⁺ signaling. 2. Ca^2+^ activates protein kinase C (PKC), which phosphorylates GluA2-containing AMPA receptors (GluA2-AMPAR), promoting their internalization and reducing AMPA-mediated transmission. 3. Concurrently, Ca²⁺/calmodulin-dependent protein kinase II (CaM-KII) is activated and reduces NMDA-mediated transmission. In addition, the trafficking of GluN2B-containing NMDA receptors (GluN2B-NMDARs) to extrasynaptic sites may be facilitated. 4. Upon glutamate release (Red dots), postsynaptic AMPA and kainate receptors (AMPAR/KAR) are activated, producing an early (synchronous) evoked excitatory postsynaptic current (eEPSC). 5. Glutamate spillover, driven by presynaptic 5-HT_2A_ receptor activation and GluN2B-NMDAR phosphorylation, is modulated by presynaptic metabotropic glutamate receptors 2/3 (mGlu 2/3) and the adenosine A1 receptor (A1). 6. The trafficking of GluN2B-NMDARs may increase the excitability of extrasynaptic sites. The diffusion of the early eEPSC to extrasynaptic regions removes the Mg²⁺ block from GluN2B-NMDAR, which are activated by glutamate spillover. They elicit a late eEPSC associated with transient up states, increasing the resting membrane potential.

The early excitatory phase is absent in LSD, due to its 5-HT_1A_ activity, and is also abolished at higher concentrations of DOB (Arvanov et al., 1999a, 1999b). This concentration and time -dependence of the early phase in drugs lacking 5-HT_1A_ agonism may reflect pharmacodynamic properties that favor presynaptic 5-HT_2A_ signaling at lower doses, like higher receptor affinity or reduced occupancy. Another explanation involves compartmentalized receptor signaling: presynaptic glutamate release may involve extracellular 5-HT_2A_ receptors, while postsynaptic inhibition may require intracellular receptor activation. At low concentrations, psychedelics could gradually diffuse across the membrane, activating intracellular receptors over time (Figures 3 and 4). This hypothesis also accounts for the prolonged effects observed after washout. Supporting this, the head twitch response, a rodent behavioral proxy for hallucinogenic activity, requires intracellular receptor activation (Vargas et al., 2023). Nevertheless, extracellular receptors appear to mediate the inhibition of action potentials, as suggested by Ekins et al. (2023), although these findings should be interpreted with caution (see Section 2.3.1). However, both reports are not mutually exclusive: it is possible that postsynaptic activity is modulated by intracellular receptors while action potentials are controlled by outer plasma membrane 5-HT_2A_. Notably, this would imply that postsynaptic activity, and not neuronal firing, is key to head twitch response.

The interplay between CaM-KII and PKC is especially intriguing. CaM-KII is a sophisticated enzyme involved in synaptic plasticity, performing key roles in both long-term potentiation (LTP) and long-term depression (LTD) (for a review, see Bayer & Giese, 2025). High levels of Ca^2+^/calmodulin promote CaM-KII binding to GluN2B-NMDARs, facilitating LTP, whereas dissociation is linked to LTD (Bayer & Giese, 2025). Interestingly, Arvanov et al. (1999a) showed that inhibiting either CaM-KII or Ca^2+^/calmodulin prevents DOB-induced suppression of NMDA-evoked currents. Additionally, Barre et al. (2016) showed that PKC enhances NMDAR conductance by phosphorylating Ser1303 in GluN2B subunit, preventing desensitization. However, this modification also disrupts CaM-KII binding (O’Leary et al., 2011). Moreover, DOI promotes internalization of GluA2-containing AMPARs through PKC, a process typically associated with LTD (Berthoux et al., 2019). Thus, suppression of LTP-related mechanisms appears to underlie desensitization to excitatory input, while LTD-associated pathways in apical dendrites maintain excitability. Adding to this paradox, rather than the spine shrinkage typically associated with LTD (Woolfrey et al., 2018), an increase in apical spine density is observed following psychedelic exposure (Shao et al., 2025). These contradictory interactions need further investigation into the roles of PKC and CaM-KII in psychedelic-induced modulation of synaptic plasticity and electrophysiology.

The modulation of GluN2B-NMDARs is remarkable in the context of psychedelic action. This subunit is unique in its sensitivity to CaM-KII (Bayer & Giese, 2025) and has been implicated in both enhanced presynaptic glutamate release (Barre et al., 2016) and the generation of late eEPSCs when activated at extrasynaptic sites (Lambe & Aghajanian, 2006). These processes are tightly coupled, as glutamate spillover from increased presynaptic release can activate these extrasynaptic receptors (Lambe & Aghajanian, 2006). The combined activity of presynaptic and extrasynaptic GluN2B-NMDARs likely contributes to the induction of transient up states (Figure 4). Moreover, GluN2B-containing receptors generate low-amplitude, slow-decay currents (Vicini et al., 1998), making them well-suited for sustaining prolonged, late eEPSCs (Lambe & Aghajanian, 2006). Additionally, their high mobility within the membrane allows them to redistribute from synaptic to extrasynaptic regions (Groc et al., 2006). Therefore, CaM-KII-dependent inhibition at postsynaptic sites may be followed by a receptor’s reorganization that enhances extrasynaptic excitability (Figure 4). This specific regulation of GluN2B-NMDARs appears central to the sustained effects of psychedelics (Davoudian et al., 2023) and warrants further study.

### 4.4 Excitatory vs Inhibitory Action of Psychedelics

*In vitro* studies show that L5p neurons highly expressing 5-HT_2A_ consistently increase their firing rates (Schmitz et al., 2022). *In vivo*, psychedelics increase the average firing of L5p neurons projecting to brainstem regions (Celada et al., 2008; Puig et al., 2003; Riga et al., 2014; Shao et al., 2025). These types of L5p cells are type I neurons (Elliott et al., 2018). Type I neurons are fewer in number, project outside the telencephalon to structures like the brainstem or the thalamus, have prominent and complex apical compartments, and respond to 5-HT with a strong 5-HT_2A_-mediated excitatory current. In contrast, type II neurons are more common, project to intratelencephalic regions, have larger basal compartments, and respond to 5-HT with more modest responses, most often inhibitory currents dependent on 5-HT_1A_ activity (Elliot et al., 2018). Therefore, psychedelics seem to specifically increase the firing of a subset of 5-HT_2A_-expressing type I neurons. These are likely the minority of cells where DOB elicited a concentration-dependent potentiation of AMPA-evoked currents (Arvanov et al., 1999a).

Consistent with the higher prevalence of type II neurons (Elliot et al., 2018), most L5p cells are inhibited by psychedelics (Wood et al., 2012). This inhibition is 5-HT_2A_-dependent and involves GABAergic transmission (Puig et al., 2003). A minority of 5-HT_2A_-expressing cells are parvalbumin-positive, fast-spiking interneurons (Weber & Andrade, 2010), in line with Dearnley et al. (2021) reporting a small subset of excited fast-spiking neurons offsetting the firing rates of a large number of inhibited cells.

However, DOI, lacking 5-HT_1A_ activity, consistently inhibits the firing of most L5p neurons when applied locally (Ashby et al., 1989; Ashby & Wang, 1990; Bambico et al., 2010; Bergqvist et al., 1999; El Mansari & Blier, 1997; Reuter & Blier, 1999). In contrast, it only excites a varying subset of L5p neurons when administered systemically (Celada et al., 2008; Puig et al., 2003; Riga et al., 2014; Shao et al., 2025). Moreover, upon local administration, L5p inhibition persists in the presence of picrotoxin and MgCl_2_ (Ashby et al., 1989; Bergqvist et al., 1999), suggesting a direct, 5-HT_2A/C_-mediated mechanism.

We have seen that psychedelics inhibit postsynaptic currents through 5-HT_2A_ receptors (Arvanov et al., 1999a; Berthoux et al., 2019). Another potential direct inhibitory mechanism of 5-HT_2A_ receptors involves the recruitment of pertussis toxin-sensitive Gɑ_i/o_ signaling by 5-HT_2A_ receptors (González-Maeso et al., 2007; Kurrasch-Orbaugh et al., 2003). Alternatively, psychedelics may hinder firing by rapidly and selectively inducing 5-HT_2A_ internalization in L5p axons, as suggested by experiments using iPSC-derived human cortical neurons (Schmidt et al., 2024). Additionally, DOI-induced inhibition could be mediated by 5-HT_2C_ receptors, which are co-expressed with 5-HT_2A_ in L5p neurons (Nocjar et al., 2015). However, knocking out the 5-HT_2C_ receptor does not prevent DOI’s inhibitory effect on L5p firing, although it intriguingly abolishes the blocking action of 5-HT_2A_ antagonism on DOI (Reuter et al., 2000). More research is needed to address this issue.

Gɑ_i/o_ recruitment by 5-HT_2A_ activity is selective to psychedelic-triggered signaling, since lisuride does not engage this pathway (González-Maeso et al., 2007). Interestingly, lisuride and BL 3912A (a non-hallucinogenic DOB analog) also fail to inhibit NMDA currents, in contrast to their hallucinogenic congeners (Arvanov et al., 1999b). Since inhibition seems to be a characteristic feature of psychedelic vs non-hallucinogenic 5-HT_2A_ agonists, the question of whether increased neural excitability truly underlies psychedelic experiences arises. For instance, optogenetic activation of neurons where DOI increases Ca^2+^ transients does not elicit the head twitch response in mice (Muir et al., 2024).

Beyond 5-HT_2A_ receptor activity, a recent K_v_7-mediated mechanism of action potential inhibition has been reported (Ekins et al., 2023; Wang et al., 2025). Although these works having certain limitations regarding their pharmacological protocols (see section *Intrinsic excitability*), this novel mechanism warrants further investigation.

Our findings show that, although 5-HT_2A_ is a canonical Gα_q/11_-coupled GPCR (Kim et al., 2020), and its activation by physiological levels of 5-HT typically produces an excitatory response (Araneda & Andrade, 1991; Arvanov et al., 1999b), psychedelics exert a generalized inhibitory effect on L5p neurons, mediated by both direct L5p modulation and GABAergic activity (Ashby et al., 1989; Bergqvist et al., 1999; Puig et al., 2003; Wood et al., 2012). Along with that inhibition, a subset of projecting L5p neurons are hyperactivated (Elliot et al., 2018; Shao et al., 2025; Schmitz et al., 2022). These neurons are in a privileged position to act like cisterns that integrate large amounts of top-down activity and send it outside the telencephalon, while those inhibited are best suited to receive bottom-up input from specific thalamic nuclei (Aru et al., 2020). Additionally, higher-order thalamocortical fibers project back to the coupling compartment of L5p neurons, producing up states that boost apical drive (Aru et al., 2020; Lambe & Aghajanian, 2006; Phillips et al., 2025). This creates a scenario where the psychedelic experience coincides with a decoupling of context-driven neurons from sensory-driven ones. This pure integrative experience fits with the feelings of understanding and revelation, often without a clear content, elicited by psychedelics in human beings (Nichols, 2016). Future research should further explore the specific implications of such functional reorganization by psychedelic drugs.

### 4.5 Relation to Functional and Behavioral Literature

Our proposal that psychedelics induce a state in which integrative, contextually driven activity takes control of cortical dynamics over specific sensory input provides a framework to reinterpret results from functional and behavioral research in the field.

For instance, fMRI studies reveal that psychedelics increase interactions among large-scale functional networks in the brain, leading to enhanced between-network and global functional connectivity (Avram et al., 2025; Barrett et al., 2020; Carhart-Harris et al., 2016; Luppi et al., 2021; Siegel et al., 2024; Tagliazucchi et al., 2016). Moreover, functional hyperconnectivity, especially in associative areas, has been shown to correlate with subjective intensity of psychedelic experiences (Mortaheb et al., 2024; Siegel et al., 2024; Tagliazucchi et al., 2016). Our scheme hypothesizes clear, experimentally grounded, and testable mechanisms underlying this key finding. ET L5p neurons possess extensive apical tufts that integrate inputs from multiple cortical columns, as well as from nonspecific thalamic projections (Aru et al., 2020). At the same time, their efferences target various subcortical regions, including higher-order thalamic nuclei and monoaminergic brainstem nuclei, which in turn project back to the cortex in a widespread manner. Consequently, selective activation of ET L5p neurons enables local cortical activity to be driven by diffuse, non-specific input, and to be broadcasted widely across the cortex via ET cortico-thalamo-cortical loops (Aru et al., 2020). This is detected as increased between-network functional connectivity in fMRI studies.

Our proposal can also be linked to behavioral studies of cognitive performance (Kaup et al., forthcoming). Emerging research is showing how psychedelics alter contextual processing in vision by modulating surround suppression (e.g., Aqil et al, 2025; Swanson et al, 2024). Moreover, context-sensitive cortical activity has been proposed as an underlying mechanism of attention, as attention depends on distinguishing objects from one another (Phillips et al., 2025; Zolnik et al., 2026). Psychedelics are known to impair performance on attention tasks in humans (for reviews about the cognitive effects of psychedelics in humans, see Bălăeţ, 2022; Basedow et al., 2024; Kase et al, 2026). We propose that psychedelics disrupt attention by inducing context hypersensitivity, which interferes with the brain’s normal ability to distinguish objects from their contextual background.

### 4.7 Limitations and Future Perspectives

The present work has several important limitations that should be acknowledged and that point toward clear directions for future research.

First, although we reviewed an extensive body of literature investigating the electrophysiological effects of psychedelics in the PFC, it remains uncertain to what extent the mechanisms described here generalize to other cortical regions. The PFC is uniquely enriched in long-range projections and associative connectivity. Its pyramidal neurons have, in general, more dendritic branches and higher spine density than their counterparts in occipital, temporal and parietal lobes (Elston, 2003). In addition, it holds the largest expression of 5-HT_2A_ receptors (Beliveau et al., 2017), which makes this region especially sensitive to psychedelic agonism (Nichols, 2016). Whether similar psychedelic-induced modulation of apical drive and context-sensitive integration occurs in other associative cortices, such as posterior parietal or temporal regions, or in primary sensory cortices, remains an open question. Addressing this limitation will require targeted in vitro and in vivo experiments across multiple cortical areas, ideally combining laminar electrophysiology with cell-type-specific manipulations.

Second, the electrophysiological evidence reviewed here derives from rodent models. While rodents provide unparalleled access to cellular and circuit-level mechanisms, important species differences exist in cortical architecture, dendritic complexity, and long-range connectivity. These differences may be especially relevant for understanding psychedelic effects, given the proposed central role of apical dendritic integration and cortico-thalamo-cortical loops. Future studies should therefore prioritize translational approaches that directly test whether key findings from rodents generalize to the human cortex.

In this regard, in vitro patch-clamp and imaging experiments in human cortical tissue obtained from neurosurgical resections (e.g., temporal or frontal lobectomies performed for therapeutic reasons) represent a critical avenue for progress. Such preparations would allow direct examination of psychedelic modulation of intrinsic excitability, dendritic integration, and synaptic dynamics in human pyramidal neurons, as well as potential species-specific differences in 5-HT_2A_ receptor distribution and function.

Nevertheless, we acknowledge that certain features of human electrophysiology cannot be recapitulated in rodent models. For example, alpha-band oscillations dominate the human occipital cortex during rest, particularly with eyes closed. Suppression of alpha power during the acute effects of psychedelics is one of the most robust findings in human EEG and MEG research (e.g., Muthukumaraswamy et al., 2013; Riba et al., 2004; Timmermann et al., 2019; Valle et al., 2016). Moreover, this reduction correlates with psychedelic-induced visual imagery and is mediated by 5-HT2A receptor activation (Valle et al., 2016), whose expression is particularly high in the human primary visual cortex (Beliveau et al., 2017). This review shows that there is no clear parallel to these oscillatory changes in rodents.

Third, a major limitation of the current framework is the lack of direct correlation between the proposed cellular mechanisms and subjective experience. While correlations between functional hyperconnectivity and ego dissolution are well established, bridging cellular-level dynamics with conscious experience remains challenging, especially in humans (Kaup et al., forthcoming). However, future work combining invasive electrophysiology, computational modeling, and high-resolution neuroimaging will be essential to test whether enhanced apical drive and context hypersensitivity are necessary and sufficient conditions for specific dimensions of psychedelic experience in animal models.

Taken together, these limitations highlight the need for a multiscale research program integrating cellular electrophysiology, circuit dynamics, systems neuroscience, neuroimaging, computational modeling and phenomenology.

## 5. Conclusion

This review reveals the complexity of the electrophysiological actions of serotonergic psychedelics on L5p neurons in the PFC. Contrary to the widespread assumption that psychedelics uniformly enhance cortical excitability, the evidence shows diverse, often opposing effects depending on dose, receptor localization, and neuronal compartment. Psychedelics influence membrane potential, synaptic transmission, firing patterns, and oscillatory dynamics. These nuanced effects cannot be fully captured by neuroimaging alone, highlighting the indispensable value of electrophysiological approaches.

Furthermore, the review supports the hypothesis that psychedelics promote apical drive in L5p neurons, potentially shifting the balance of cortical processing toward top-down influence. This has significant implications for our understanding of psychedelic-induced altered states of consciousness and challenges prevailing models. Moving forward, integrative studies combining electrophysiology, imaging, and computational modeling will be essential to unravel the multi-scale mechanisms underlying psychedelic effects on brain function.

## Acknowledgements

Graphical Abstract: Created in BioRender. Jiménez, J. (2026) https://BioRender.com/v78d6y0

Figure 1: Created in PRISMA flow diagram generator (Haddaway et al., 2022)

Figure 2: Created in BioRender. Jiménez, J. (2025) https://BioRender.com/8tc7nxw

Figure 3: Created in BioRender. Jiménez, J. (2025) https://BioRender.com/6xk0pkn

Figure 4: Created in BioRender. Jiménez, J. (2025) https://BioRender.com/ds00zvg

